# DNACSE: Enhancing Genomic LLMs with Contrastive Learning for DNA Barcode Identification ^†^

**DOI:** 10.1101/2025.10.27.684901

**Authors:** Jiadong Wang, Bin Wang, Shihua Zhou, Ben Cao, Wei Li, Pan Zheng

## Abstract

DNA barcoding is a powerful tool for exploring biodiversity, and DNA language models have significantly facilitated its construction and identification. However, since DNA barcodes come from a specific region of mitochondrial DNA and there are structural differences between DNA barcodes and reference genomes used to train existing DNA language models, it is difficult to directly apply the existing DNA language models to the DNA barcoding task. To address this, this paper introduce DNACSE (DNA Contrastive Learning for Sequence Embeddings), an unsupervised noise contrastive learning framework designed to fine-tune the DNA language foundation model while enhancing the distribution of the embedding space. The results demonstrate that DNACSE outperforms direct usage of DNA language models in DNA barcoding-related tasks. Specifically, in fine-tuning and linear probe tasks, it achieves accuracy rates of 99.17% and 98.31%, respectively, surpassing the current state-of-the-art BarcodeBERT by 6.44% and 6.44%. In zero-shot clustering tasks, it raises the Adjusted Mutual Information (AMI) score to 92.25%, an improvement of 8.36%. In addition, zero-shot benchmarking and genomic benchmarking tests are evaluated, indicating that DNACSE enhances the performance of DNA language models in generalized genomic tasks. In summary, DNACSE has demonstrated excellent performance in DNA barcode species classification by making full use of multi-species information and DNA barcode information, providing a feasible way to further explore and protect biodiversity. The code repository is available at https://github.com/Kavicy/DNACSE.

## Introduction

DNA barcodes are short DNA fragments from specific genes used as a critical tool for assessing biodiversity.^1^ They help identify and characterize organisms and are vital in fields such as ecology^2,3^, food safety^4^, and public health.^5^

Different organisms use different genetic markers for DNA barcoding. The mitochondrial Cytochrome c Oxidase I (COI) gene is the standard for animals and some protists,^6^ and the internal transcribed spacer (ITS) region is widely used in fungi. ^7^ However, a universal barcoding region for plants has yet to be identified. Consequently, a combination of markers, such as internal transcribed spacers (ITS1 and ITS2) and chloroplast genes (matK and rbcL), are commonly used for plant taxonomic identification. ^8^

To utilize these gene fragments for species identification, researchers have developed multiple methods. Generally, species classification methods using DNA barcodes can be broadly categorized into two main types. The first type relies on alignment-based approaches, such as BLAST,^9^ which determine the taxonomic category of a sequence by aligning the query sequence against reference sequences in a database. However, these methods struggle to meet practical demands when processing massive datasets due to their high computational costs and time-consuming nature. The second type of approach consists of prediction-based methods, which utilize techniques such as k-mer frequency or deep learning for classification. Although these approaches are more efficient, they generally suffer from insufficient generalization capabilities and fail to effectively comprehend the deeper semantic information within DNA sequences. A variety of models and methods have been developed using these approaches to address specific taxonomic challenges. For example, AIfie^10^ systematically evaluated five algorithms: K-Nearest Neighbors (KNNs), Support Vector Machines (SVMs), Random Forests (RFs), Extreme Gradient Boosting (XGBs) and Deep Neural Networks (DNNs) and performed kingdom-level classification on the animal COI gene dataset. The CNN_FunBar^11^ model employs a Convolutional Neural Network (CNN) to classify fungi at different taxonomic levels. The ESK^12^ model combines an Elastic Network Stacked Autoencoder (EN-SAE) with Kernel Density Estimation (KDE) to classify fish across different families.

To address these challenges, a new paradigm has emerged: the DNA language model. These models significantly facilitate species classification by first performing unsupervised pre-training on a wide range of DNA sequences, followed by a fine-tuning paradigm to migrate the learned representations to downstream genomic tasks. Within this paradigm, two primary strategies for pre-training data have been adopted, each with its own set of challenges. First, some models use large-scale human reference genomes or other taxonomically irrelevant genomic regions for pre-training.^13–15^ However, the distribution differences between these general data and specific DNA barcodes lead to a domain shift problem, ^16^ which impairs the performance in barcoding tasks. Second, another approach is to pretrain models from scratch exclusively on DNA barcodes. BarcodeBERT,^16^ for example, was the first model trained specifically for invertebrate DNA barcoding and has shown superior overall performance in both closed-world and open-world settings. Although it has set the current benchmark, subsequent models like BarcodeMAE^17^ and Barcodemamba^18^ have improved performance in specific tasks such as barcode index number (BIN) reconstruction and the linear probe task. Despite these advancements, these models are computationally expensive to train and, due to their limited training data, they lack the broader biological context needed for effective generalization. Beyond these specific challenges, a shared limitation affects both types of DNA language models: the anisotropy of transformer embedding spaces.^19^ This issue, common in models trained with likelihood loss, causes word vectors to become clustered in a narrow, cone-shaped region of the embedding space. This can significantly impair model performance on downstream semantic similarity tasks, such as predicting taxonomic labels.

To address the aforementioned challenges, this paper proposes an unsupervised contrastive learning framework named DNACSE. It leverages DNA barcodes to perform domain adaptation on pre-trained DNA language models. Unlike training a new model from scratch, our approach aims to fine-tune existing models to better suit DNA barcoding tasks. This achieves outstanding performance while significantly reducing training costs.

The core idea of DNACSE is to combine the rich contextual semantic representations of traditional DNA language models with domain knowledge from DNA barcodes, thereby effectively addressing the domain shift problem faced by multi-species DNA language models. Specifically, this paper designed a carefully constructed feature augmentation layer that generates diverse positive sample representations by introducing perturbations at the feature level, thereby simulating unexpected scenarios during the sequencing process. Simultaneously, to mitigate the interference of randomly generated negative samples during training, this paper introduced a hard negative sample construction module. ^20,21^ This module generates samples that are similar, yet distinct from positive samples, thereby enhancing the model’s discriminative capability.

Experimental results demonstrate that DNACSE leverages rich multi-species background knowledge to better understand DNA barcode features, outperforming baseline DNA language models across multiple tasks in both closed-world and open-world settings. In particular, in species-level supervised fine-tuning tasks, DNACSE achieves an accuracy of 99.17%, surpassing the current state-of-the-art BarcodeBERT model. In the linear probe task, the accuracy reached 98.31%, representing a 6.44% improvement over BarcodeBERT. For the BIN reconstruction task, the adjusted mutual information (AMI) ^22^ increased to 92.25%, showing an 8.36% gain compared to BarcodeBERT. Furthermore, in the Gene-MTEB^23^ and Genomic Benchmark^24^ evaluations, DNACSE achieved state-of-the-art performance in 7 and 5 tasks, respectively, demonstrating superior generalization capabilities. Collectively, these results validate that DNACSE effectively optimizes the embedding space of DNA language models, significantly enhancing their performance across downstream tasks.

## Materials and Methods

Following contrastive learning proposed by Gao et al.^25^ DNACSE fine-tunes DNABERT-2^26^ with a contrastive objective to optimize the spatial distribution of DNA embeddings and mitigate the domain shift problem observed on DNA barcodes. Specifically, for each DNA sequence, two independent forward passes through the encoder are performed to obtain different vector representations, which constitute a positive pair, while the representations of other sequences within the same batch are treated as negatives. All parameters of DNABERT-2 are updated during fine-tuning to maximize the similarity of positive pairs and minimize the similarity to negatives.

To further strengthen the learning process, several additional strategies are incorporated. First, this paper introduce Gaussian feature noise to the hidden layer to perturb the representations of positive sample pairs. This method simulates unexpected variations from sequencing, thereby enhancing the model’s robustness and enabling it to learn stable embeddings. Unlike the dropout noise used in SimCSE,^25^ DNACSE employs an additive Gaussian perturbation. This noise is intentionally applied from both global and local perspectives to the original embeddings. Second, to address the limited diversity of negatives from within-batch sampling, this paper adopts the hard negative construction strategy proposed in MixCSE,^21^ which extends SimCSE by explicitly generating challenging negatives. In DNACSE, this strategy is adapted to DNA barcodes by mixing the feature representations of positive and negative samples, thereby constructing hard negatives that partially carry positive features. These mixed negatives are deliberately confounding and force the model to learn more discriminative representations. Third, to optimize the model’s performance on contrastive learning, this paper added a nonlinear projection head. This head, composed of two linear layers and a Gaussian Error Linear Unit (GELU)^27^ activation function, projects the encoder’s output into a dedicated contrastive learning space. By doing so, it helps the model improve its invariance to nonlinear variations, a benefit consistently demonstrated in both vision^28^ and nature language processing (NLP).^25^ The final contrastive loss is then computed in this projected space, with the resulting gradients used to backpropagate and optimize the model.

The overall workflow of DNACSE, including sample construction, noise augmentation, hard negative generation, and projection into the contrastive space, is summarized in Figure 1 and Algorithm 1.

**Figure 1.**
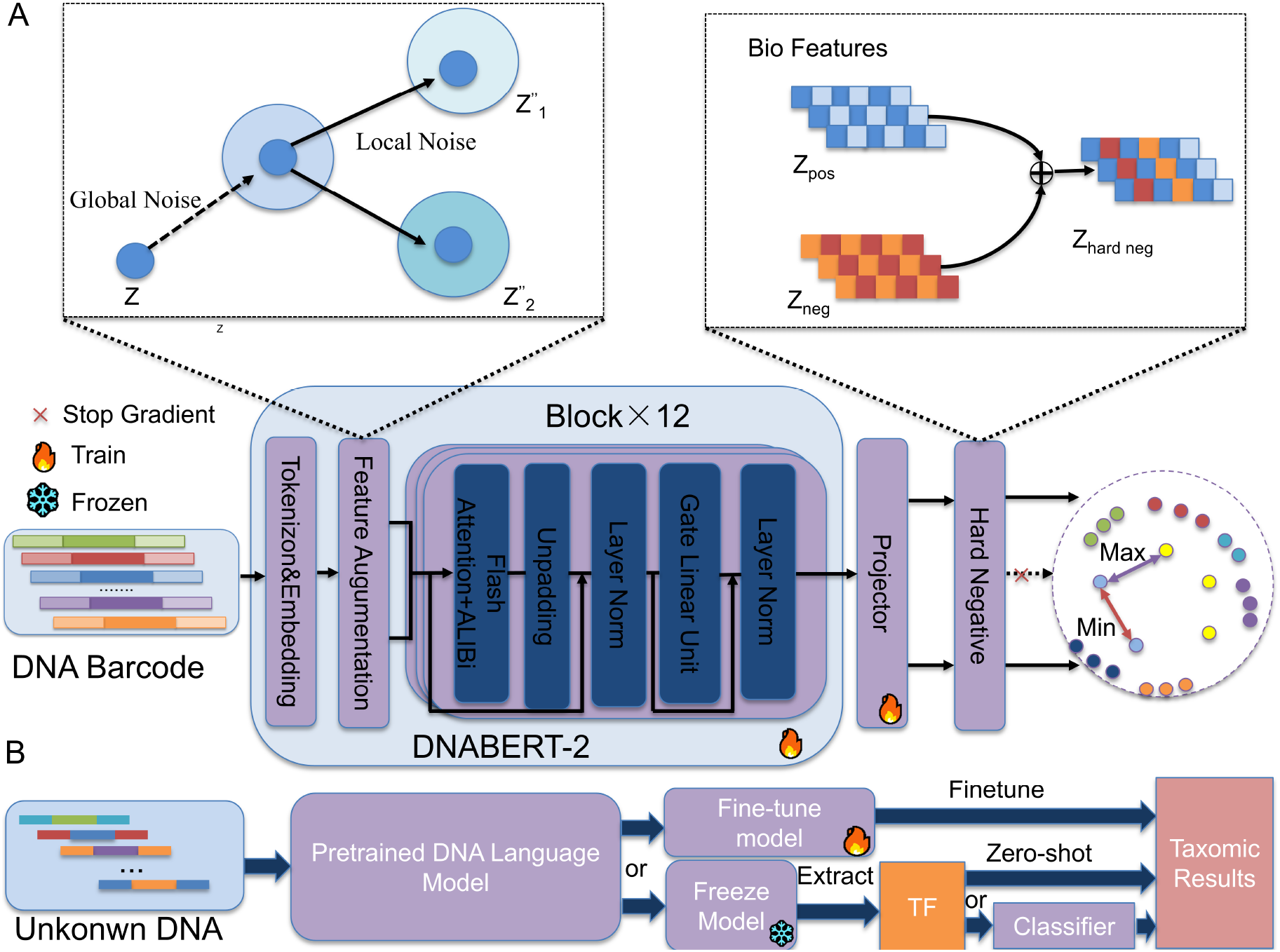
DNACSE architecture and downstream task pipeline. (A) In the contrastive learning stage, DNACSE incorporates feature augmentation and hard negative sample construction into unsupervised contrastive learning, enabling the model to learn comprehensive DNA barcode representations and capture their internal relationships. All backbone parameters are trainable in this stage, with no frozen components. (B) For downstream applications, DNACSE offers three modes of transfer. In the first mode, the entire model is fine-tuned with labeled data for task-specific adaptation. Alternatively, the pretrained backbone can be frozen to extract task-specific features, which are either used to train a lightweight classifier or directly applied in a zero-shot setting.These complementary strategies enable DNACSE to address a broad spectrum of DNA barcoding tasks with varying amounts of supervision. **Notation:** *Z* denotes the original sequence embedding, *Z*^*′*^ the global noise-augmented embeddings, *Z*^*′′*^ the final-augmented embeddings, *Z*_pos_ the positive embedding, *Z*_neg_ the negative embedding, and *Z*_hard neg_ the mixed hard negative embedding.

### Data Processing

The training dataset used in this paper is from the Bioscan-5M dataset, ^29^ a large-scale multimodal dataset containing 2.4 M unique DNA barcodes, divided into four main sections:

i. *Pretraining* : contains unique DNA barcodes with the number of 2.28 M from unclassified samples for unsupervised pre-training.
ii. *Seen*: consists of DNA barcodes with confirmed scientific species names, organized into training subsets (118,000 barcodes), validation subsets (6,600 barcodes), and test subsets (18,400 barcodes) for closed-world classification tasks.
iii. *Unseen*: comprises novel species with provisional taxonomic labels, split into subsets of reference (12,200 barcodes), validation (2,400 barcodes), and test (3,400 barcodes) subsets for open-world species identification tasks.
iv. *Other* : species containing species that do not occur in any of the other partitions form a separate subset.

Notably, each sample in the unseen partition shares only its genus-level classification with the seen partition, ensuring no species-level overlap occurs.^30^ For unsupervised pretraining, this paper collected more than 2.47M DNA barcodes from both the pretraining set and other sources, followed by a series of preprocessing steps: discarding trailing unknown bases (N), uniformly truncating sequences to 660 bp, and removing the shortest 5% of sequences. These operations yield a final dataset of approximately 2.2M unique barcodes with lengths between 640–660 bp. Importantly, truncating sequences to 660 bp and filtering out the shortest 5% are intended to maintain relative consistency in sequence length during contrastive learning. By doing so, this paper aims to reduce spurious signals arising purely from length differences, improving the consistency of our contrastive learning embeddings.

### Feature Augmentation Layer

Simple dropout is insufficient to generate meaningful perturbations in the hidden embeddings of DNA barcodes, which often results in poor generalization. To address this, DNACSE adopts additional feature augmentation as an implicit data augmentation strategy, directly perturbing hidden representations rather than raw sequences. This approach creates multiple semantically consistent views of each sequence by applying feature-level transformations, thereby encouraging the model to learn view-invariant embeddings. It also increases the diversity of the training data distribution, which is crucial for generalization and robustness in contrastive learning.^31,32^ In contrast, sequence-level operations such as random insertions, deletions, or substitutions may destroy the positional structure of nucleotides and thus harm biological plausibility. ^33^

To implement feature augmentation, DNACSE introduce a feature noise layer inserted between the hidden layers of the encoder. This layer combines two complementary perturbation mechanisms: a global noise module and a local noise module. First, the global noise module applies an additive Gaussian perturbation that is consistent across all views of the same sequence but independent across different sequences. Formally, given an input embedding *E* ∈ ℝ^*B×N ×L×H*^, where *B* is the batch size, *N* are the number of views, *L* is the sequence length, and *H* is the hidden dimension, this paper generates a global noise tensor 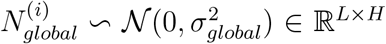. This perturbation enhances consistency within positive pairs while amplifying inter-sample variation. Second, the local noise module applies Gaussian perturbations selectively to token positions according to a predefined mutation rate, thereby simulating random sequencing errors or natural variability. The resulting local noise 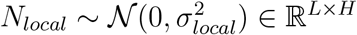 is independently generated for each view. The final augmented embedding is expressed as:

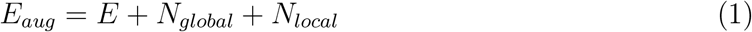

where *E*_*aug*_ ∈ ℝ^*B×N ×L×H*^. This design ensures that perturbations are subtle yet diverse, capturing both systematic and stochastic sources of variability in DNA sequencing. The feature noise layer applied only in the training phase.

### Mixed Negative Samples

Currently, training set contains a large number of DNA sequences from species with unique characteristics.^34^ These sequences provide clear guidance for model optimization during the early stages of training. However, as training progresses, the model becomes prone to overfitting. To address this, DNACSE continuously challenges the model by constructing more challenging hard negative examples.

Contrastive learning maximizes the lower bound of mutual information between an anchor and its positive view by comparing the anchor with multiple negative samples. Typically, a higher number and quality of negative samples leads to better model performance.^35^ However, negative samples randomly sampled within batches are often of insufficient quality. They may differ significantly from positive samples, failing to provide stable training motivation,^20,21^ or they may pose a risk of being pseudo-negative samples, which mislead the model into incorrectly classifying positive samples as negative.^36^

By manually introducing hard negative samples that are semantically similar to positive samples but belong to a different category, DNACSE can construct a collection of negative samples with consistent quality. This approach not only solves the problem of unstable negative sample quality, but also forces the model to learn more nuanced and subtle distinctions between semantically similar, yet distinct examples.

Given that DNA barcodes have minimal intra-species differences and significant differences between distantly related species. In some closely related species or complexes, the differences between species are relatively unclarified, resulting in unclear boundaries between species.^37–39^ In addition, each base of a DNA barcode can be critical for species identification. Random base changes can easily make a sequence non-compliant or biologically meaningless, turning it into a simple negative sample rather than a sequence that is subtly different from a positive sample but from a different species, which is what we would want.

To overcome this, DNACSE follows the feature-level mixing strategy recently introduced in NLP.^21^ Specifically, instead of perturbing sequences directly, DNACSE mix the feature representations of positive and negative samples to synthesize hard negatives that partially inherit positive features while remaining distinct. This approach, inspired by MixCSE, avoids the biological uncertainty of sequence-level perturbations and creates deliberately confounding negatives that encourage the model to learn more discriminative and robust representations in the DNA feature space.

For encoder *f*_*θ*_ and anchor sample *x*_anchor_, we mix its positive sample *x*_pos_ and any negative sample *x*_neg_ within the batch by the mixing coefficient *λ* to construct a negative sample representation 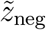, which carries the characteristics of the positive sample and is more challenging to construct. The constructive expression is as follows:

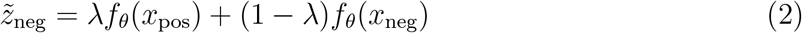

where *λ* stands for the mixing control hyperparameter. For positive samples of anchor samples, we can use the same construction method to obtain its hard negative sample. The range of *λ* is between [0, 1]. When *λ* is larger, the constructed hard negative samples are closer to the positive sample features. However, we still need to avoid constructing hard negative samples that are closer to the anchor than the positive samples to avoid unnecessary confusion. Note that these synthesized pseudo-negative samples will not participate in the gradient update process. This prevents the model from erroneously pushing the positive samples away from the anchor samples. By including positive samples with all hard negative samples, DNACSE obtains a contrastive learning loss representation that includes hard negative samples.

### Loss Function

To learn an effective representation for DNA barcodes, DNACSE adopts a modified Information Noise-Contrastive Estimation (InfoNCE) loss ^40^ as its training objective. This loss function is designed to optimize the model such that embeddings of the same species are clustered compactly, while embeddings of different species are kept well separated. The ultimate goal of this optimization is to produce a model representation space that is uniformly distributed and effective for DNA barcoding tasks.

The InfoNCE loss leverages the structural information within both positive and negative samples to generate supervised signals without relying on external labels. Therefore, the training objective focuses on minimizing the distance between positive sample pairs while maximizing the distance between negative sample pairs. In this paper, we obtain two perspectives for our training objective by applying a combination of the model’s own dropout and feature noise to the input sequences. This ensures that the model learns both denoising and alignment simultaneously.

To further increase training difficulty and prevent over alignment between positive samples, which can lead to representation collapse. We incorporate an adversarial perturbation by subtracting a fixed value from the pairwise similarity between the anchor and its positive sample. This approach aligns with the goal of spatial semantic interpolation and forces the model to explore subtle boundaries within the feature space.^41–43^ The contrastive learning goal of the DNACSE can be summarized in the following form:

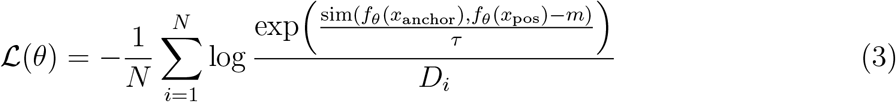

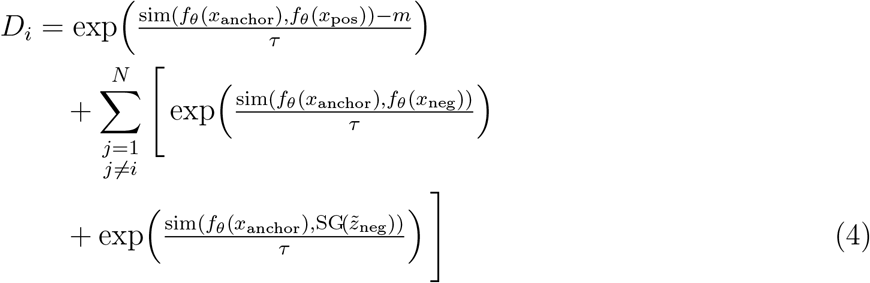

where 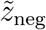 is the hard negative representation defined by formula 2. *f*_*θ*_ is a parameterized DNA language model encoder, *x*_anchor_ and *x*_pos_ is the anchor point and the positive sample

#### Algorithm 1

Learning algorithm from DNACSE.

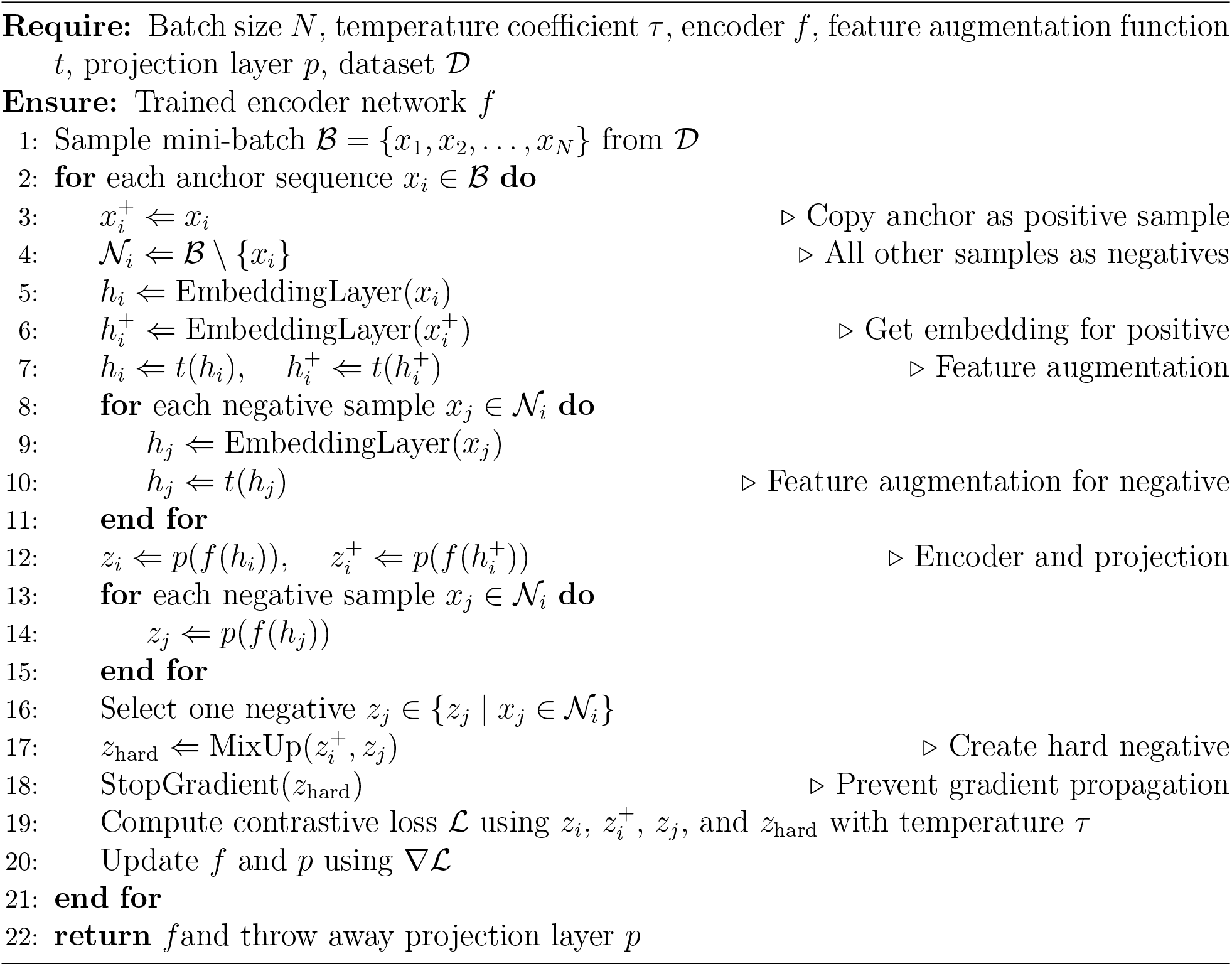

view of the *i* th sequence, *j* denotes the index of other potential negative samples within the batch, used to compute the similarity contribution between the *x*_anchor_ and other negative samples. *sim*(·) represents the cosine function, *m* is the positive sample similarity penalty, *τ* is a temperature parameter used to control smoothing of the similarity distribution, and *N* is the batch size. *SG*(·) denotes the stop gradient operator,^44^ ensuring that the hard negative samples are not updated via gradient propagation, consistent with prior work.^21^ This is a simple and effective method that can capture the semantic similarity between DNA sequences.

### Experiment setup

The DNACSE model is based on the BERT ^45^model architecture, and Flash Attention 2^46^ is used to obtain a more efficient attention representation to improve inference speed and reduce memory usage. Attention with Linear Biases (ALiBi) ^34^ is used to broaden the context length, and to capture more complex expression patterns of DNA barcode data and improve model performance, we choose to replace the Gated Exponential Linear Unit (GEGLU)^47^ activation function with the Swish-Gated Linear Unit (SwiGLU)^47^ activation function that has fewer parameters and is more efficient.

#### Flash Attention with ALiBi

For transformer-based foundation models, the run-time and memory requirements of attention layers increase quadratically with sequence length, which is the primary bottleneck when scaling them to longer sequences. This limitation, coupled with restricted context length and poor generalization capabilities, often renders them incapable of addressing the challenges posed by long genomic sequences. To address this issue, DNACSE aligns with DNA language models such as DNABERT-2 and DNAGRinder^48^ by optimizing memory usage during training and inference through a fast attention^49^ mechanism that fully leverages hardware acceleration. DNACSE employs relative position encoding techniques to enable the model to extend to context lengths not encountered during training during inference. Notably, unlike the Flash Attention mechanism in DNABERT-2, which focuses on long sequence use cases, the Flash Attention 2 mechanism in DNACSE achieves nearly double the acceleration through a series of improvements while further optimizing inference capabilities on short sequences. Since most DNA barcodes are shorter than 1,000 bp, this improvement is particularly beneficial for DNA barcoding tasks. The attention calculation process is as follows: for a query vector **q**_*i*_, key matrix **K**, value matrix **V**, and sequence length *n*, the attention output for head *i* is given by:

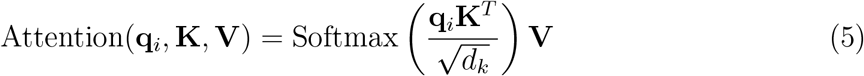

where *d*_*k*_ is the dimension of the key vectors, and the softmax is applied over the sequence length *n*. Flash Attention 2 optimizes this computation by tiling the matrices and minimizing memory transfers between Static Random Access Memory (SRAM) and High Bandwidth Memory (HBM), ensuring efficient processing even for large sequences. Additionally, FlashAttention 2 significantly accelerates attention computation through a series of kernel-level optimizations. It reduces non-matrix multiplication (non-MatMul) floating-point operations and converts as many computations as possible into efficient matrix multiplication operations. Simultaneously, it parallelizes forward and backward propagation across the sequence length dimension and performs finer-grained task decomposition. Even within a single attention block, it distributes work across distinct thread blocks to minimize communication and shared memory read/write operations. This maximizes GPU parallel processing capabilities while reducing memory I/O, ultimately achieving substantial acceleration. In order to further improve the DNA language model’s ability to generalize to longer sequences, DNACSE chose to use a simpler and more efficient extension method: ALiBi. By simply adding a linear distance penalty term to the attention score, it enables the model to perceive the position of tokens, thereby improving the model’s generalization ability during inference. The flash attention calculation with linear offset is as follows:

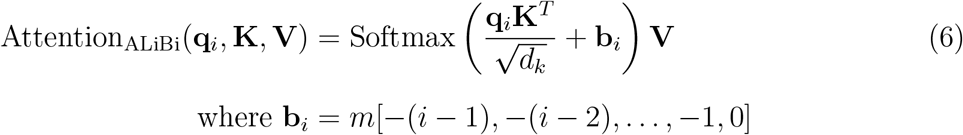

where the bias term *m*[−(*i*− 1), −(*i*− 2), …, −1, 0] is an arithmetic sequence with a common difference of −*m*, applied to the attention scores before the softmax operation. This bias reduces the attention scores for tokens further apart, enhancing the model’s ability to handle long sequences.

#### SwiGLU

To improve computational efficiency and further increase inference speed, DNACSE replaced GEGLU with SwiGLU, a more efficient variant of Gated Linear Units (GLU).^50^ SwiGLU combines a gating mechanism with the Swish activation function, which not only reduces the number of model parameters, but also captures complex data expression patterns of DNA barcodes through a dynamic gating mechanism. The SwiGLU calculation process is as follows: for an input vector **x**, weight matrices **W** and **V**, the SwiGLU output is calculated as follows:

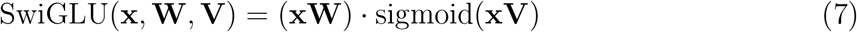

where:**x** ∈ ℝ^*d*^ is the input vector, **W, V** ∈ ℝ^*d×d*^*′* are learnable weight matrices, sigmoid 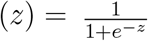 is the sigmoid function, · denotes element-wise multiplication.

DNACSE makes full use of the rich multi-species semantic information of DNABERT-2 as a basis for utilizing contrastive learning to address the poor performance of traditional DNA language models on DNA barcodes. We completed model pre-training on two RTX3090, and all downstream evaluation tasks were done on one RTX3090. The batch size allocated in each device was 64. Training was carried out for a total of one epoch with a learning rate of 3e-5 with the AdamW optimizer.^51^ A linear scheduling strategy with a warm up ratio of 0.1 was used. 30 rounds of hyper-parameter search were performed using optuna^52^ with a final choice of 0.07 for temperature *τ*, 0.02 for the positive sample similarity penalty, 0.04 for the global noise intensity, a local noise intensity of 0.15, and a variability of 0.10. We compare DNACSE to previous optimal methods on the DNA barcoding task, including three encoder-only Transformer models: DNABERT-2, DNABERT-S,^33^ and Nucleotide Transformer-multispecies-v2-50m (NT),^53^ a state-space model: HyenaDNA-tiny-1k (Hyena).^54^ The models mentioned above were pre-trained in non-DNA barcode, and BarcodeBERT^16^ pre-trained on DNA barcode.

#### Evaluation Metrics

We used a set of standard classification and clustering metrics to evaluate DNACSE performance against various baselines.^23,29^ These metrics provide a comprehensive overview of the model’s ability to correctly classify and cluster DNA barcodes from different species, as well as its generalization ability in broader genomic tasks. The following metrics were calculated:

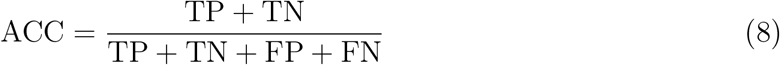

where true positives (TP) is the number of samples correctly predicted as positive, true negatives (TN) is the number of samples correctly predicted as negative, false positives (FP) is the number of samples incorrectly predicted as positive, and false negatives (FN) is the number of samples incorrectly predicted as negative.

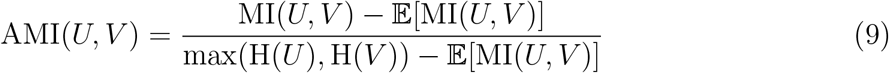

where 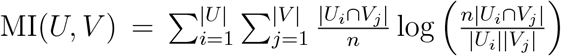 is the mutual information between the clustering results *U* and the true classification 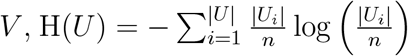 is the entropy of the clustering results *U*, H(*V*) is the entropy of the true classification *V, n* is the total number of samples. 𝔼[MI(*U, V*)] is the expected mutual information.

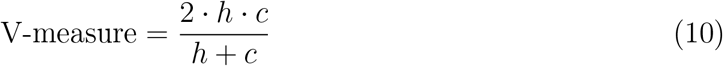

where 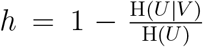 is the homogeneity 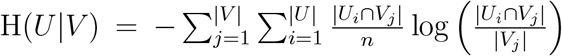 is the conditional entropy of true classification *U* given the clustering results 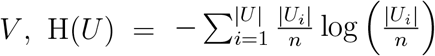 is the entropy of true classification *U, n* is the total number of samples.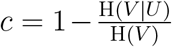 is the completeness, H(*V* |*U*) is the conditional entropy of the clustering results *V* given the true classification *U*, H(*V*) is the entropy of the clustering results *V*.

## Results

To give a fair overview of how DNACSE performed compared to other DNA language models in a variety of genomic tasks. First, we test DNACSE on five DNA barcoding tasks, including three closed-world tasks and two open-world tasks. Next, we use a zero-shot benchmark (MTEB-Gene^23^) and a fine-tuning benchmark (Genomic Benchmark^24^) to demonstrate that DNACSE can improve the distribution of DNA language model embeddings, thus enhancing performance in downstream tasks. Investigate the influence of each DNACSE module on the zero-shot clustering task of DNA barcoding and determine which data augmentation methods are more suitable for DNA barcoding data. Finally, we carried out a series of ablation experiments to explore and explain the different effects of various configurations of DNACSE and different data augmentation methods.

### DNA Barcoding Tasks

This paper uses different self-supervised learning objectives to compare the performance of DNACSE with other baseline models to assign DNA barcodes to the corresponding taxonomic classes. In a closed-world setting, these tasks are designed to assign DNA barcodes to known taxonomic units, encompassing three distinct subtasks. In the fine-tuning task, the pre-trained model is trained on a training subset of the *seen* partition and evaluated on a test subset, which is used to test the model’s performance on a species-specific classification using exclusively labeled data. In the linear probe and 1 Nearest Neighbor (1NN) probe tasks, we evaluate the quality of the pre-trained embeddings by freezing the model’s backbone. Specifically, in the linear probe task, we utilize model embeddings to train a linear classifier on the *seen* partition and evaluate its performance on a test subset. In the 1NN probe task, we evaluate the model using *unseen* partitions as the query set and the training subset of known partitions as the reference set. The probe is conducted by identifying the nearest neighbor on the basis of cosine similarity. The goal is to assign the barcodes in the query set to the nearest category labels in the training subset at the genus level, and this task is used to evaluate the model’s ability to generalize to new species in known genera. All the results of the above experiment are evaluated using accuracy as a metric, with higher accuracy representing higher performance. In an open-world setting, these tasks are to cluster sequences from unknown species based on shared features. Specifically, we used BIN and species categories as clustering labels and set up a BIN reconstruction task and a species-level clustering task. we freeze the model backbone, mix the test subset of the *seen* partition with the test subset of the *unseen* partition, and use an encoder to obtain their embeddings. The embeddings were captured using a model and down-scaled to 50 dimensions with UMAP.^55^ The agglomerative clustering^56^ was then applied to these embeddings (using the L2 distance and Ward’s method). The clustering result was used to assess the model’s ability to capture the hierarchical structures of taxonomic relationships for rare or unclassified species. All the above results were evaluated using AMI as an evaluation metric, with higher scores representing better clustering results.

The results in Table 1 and Figure 2 demonstrate that the outstanding performance of DNACSE is due to two key factors. First, the rich multi-species knowledge of its pre-trained model provides DNACSE with more generalized and superior initial embeddings compared to BarcodeBERT, which was pretrained exclusively on DNA barcodes. This gives DNACSE an advantage in using generic genomic features to understand DNA barcodes. Second, by performing rapid fine-tuning on DNA barcodes using unsupervised contrastive learning, DNACSE quickly adapts to the structural information of DNA barcodes, which is another crucial reason for its success on DNA barcoding tasks, outperforming models pretrained on non-barcode. The final results show that DNACSE exceeds all comparative baselines, achieving the best classification accuracy on fine-tuning and linear probe tasks, as well as the highest AMI scores on the BIN reconstruction task and the species-level zero-shot clustering task. Specifically, on the linear probe task, DNACSE outperforms suboptimal DNABERT-S by 2.82% and showed a 30.55% improvement over DNABERT-2. In addition, it also shows a 6.44% improvement compared to the pre-trained form scratch BarcodeBERT. Furthermore, DNACSE achieved the best results in the species-level zero-shot clustering task, improving by 0.34% over second-place DNABERT-S and by 6.79% over BarcodeBERT. These findings collectively demonstrate that combining the rich multi-species background knowledge from a pre-trained DNA language model with DNA barcodes successfully enhances the model’s performance on the DNA barcoding task.

**Table 1:**
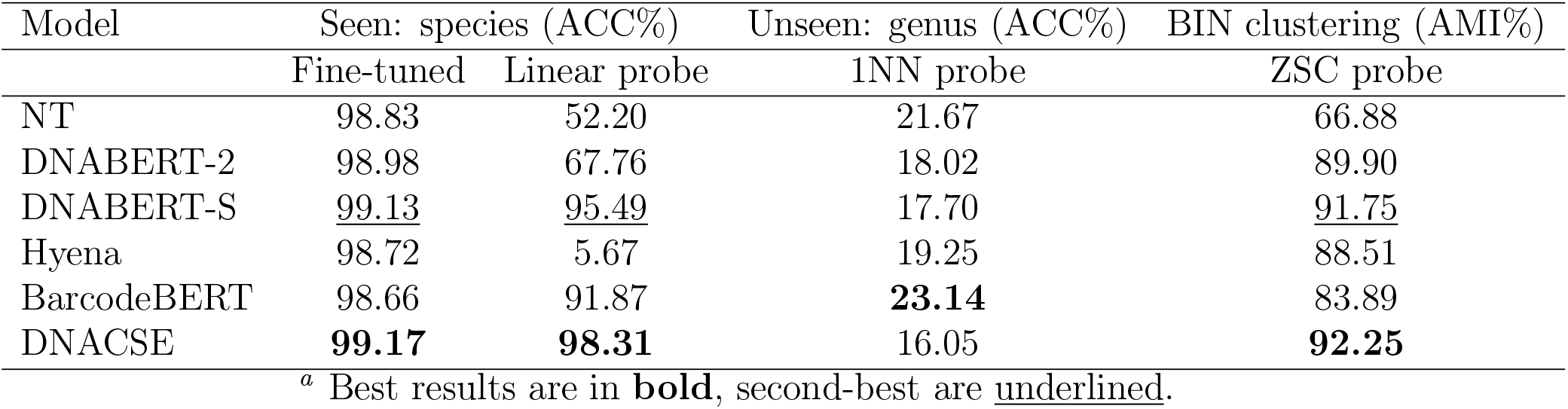
Performance metrics of different models on DNA barcoding tasks^*a*^.

**Figure 2.**
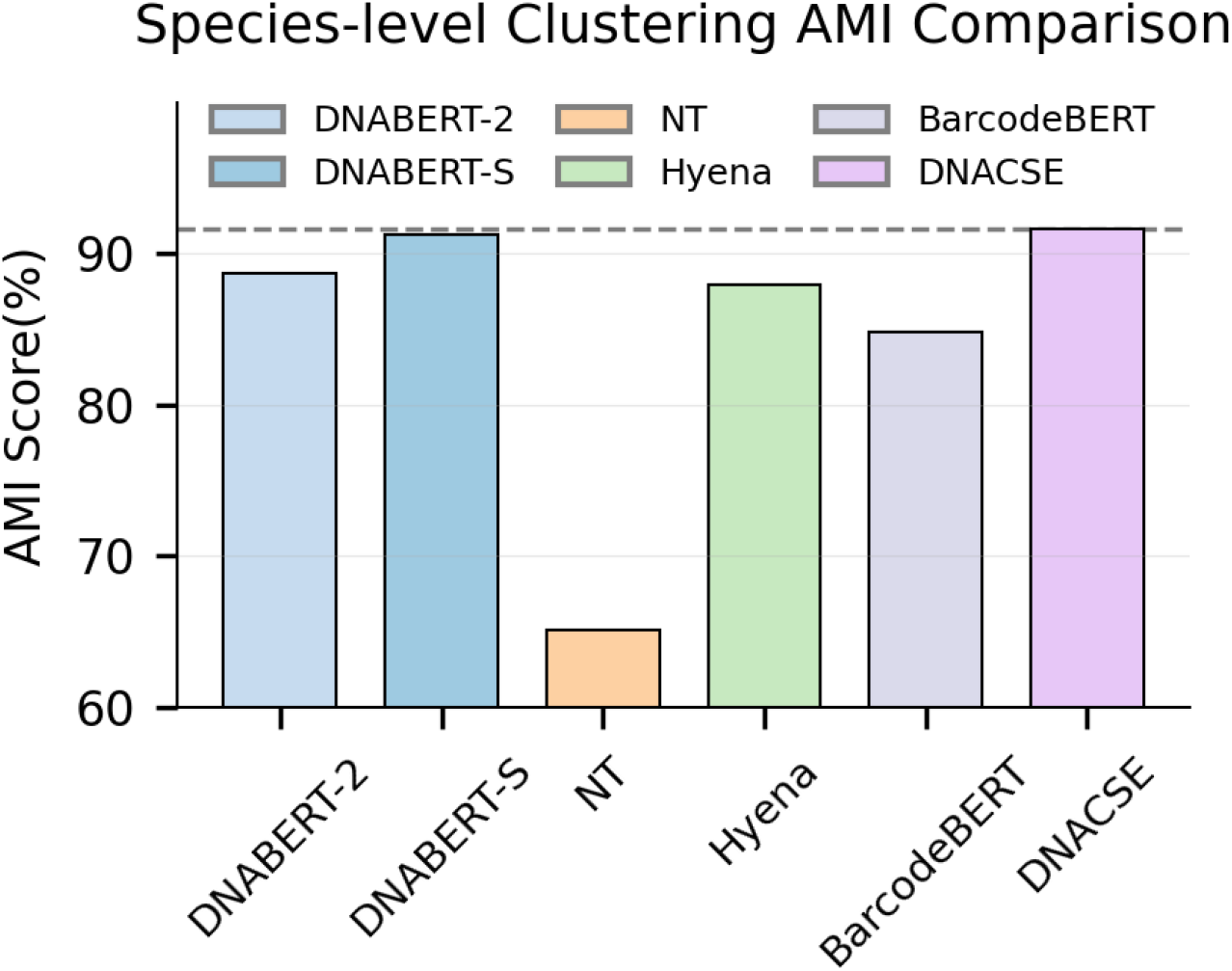
Species-level zero-shot clustering AMI (%) performance

While DNACSE achieves clear improvements in fine-tuning, linear probing, and zeroshot clustering tasks, its performance on the 1-NN probe is relatively weaker. This discrepancy can be explained by the different sensitivities of these evaluation protocols. The 1-NN probe relies exclusively on raw embedding distances, making it highly sensitive to local decision boundaries and rare taxonomic signals.^57^ In contrast, fine-tuning and linear probing exploit global discriminative features, which are better aligned with the design of DNACSE. DNACSE introduces feature noise, hard negative construction, and a contrastive learning objective with a positive-sample margin penalty. These components explicitly enhance global separability in the embedding space, leading to strong performance on tasks requiring global discriminability,^25^ but at the cost of reduced locality preservation. By comparison, models pretrained with masked language modeling (e.g., BarcodeBERT) tend to maintain fine-grained local similarities due to the context-recovery nature of their objective, which naturally benefits 1-NN performance but offers weaker global clustering structure.^58^ This contrast highlights a fundamental trade-off between global discriminability and local neighbor preservation^59^ in DNA barcode embeddings.

To visually assess the quality of each model embedding, this paper performed a two-dimensional visualization analysis of embeddings from multiple DNA language models, including DNACSE, DNABERT-2, and DNABERT-S, on the BIN reconstruction dataset. We employed UMAP to project the high-dimensional embedding vectors onto a two-dimensional plane for intuitive assessment of their intrinsic structure. Each point’s color represents distinct BIN labels. As shown in Figure 3, by comparing the results, this paper observed that DNACSE demonstrates exceptional local clustering capability: the UMAP plot clearly reveals that the embeddings generated by DNACSE exhibit the strongest structural coherence. Although achieving global separation is extremely challenging under zero-shot and massive-category settings, sequences from the same taxonomic unit tend to cluster together locally, forming numerous compact small clusters. This visually demonstrates that DNACSE successfully learned feature representations with high discriminative power that reflect biological taxonomic relationships. In contrast, the embedding space of DNABERT-S exhibits significant flaws, with a pronounced “void” appearing in the central region of its UMAP plot. This indicates an incomplete or biased learned manifold structure, suggesting the model fails to effectively utilize the entire representation space. The DNABERT-2 embedding space presents as a highly mixed scatter cloud. Data points of the same color are randomly scattered throughout the space, indicating that the learned features possess very limited ability to distinguish between different species. In summary, these UMAP visualizations provide compelling visual evidence for our conclusions. The embedding space learned by DNACSE is not only structurally superior and more continuous but also semantically meaningful, mapping biologically similar sequences to closer positions. This high-quality representation learning capability is the fundamental reason behind DNACSE’s leading performance in the BIN reconstruction task.

**Figure 3.**
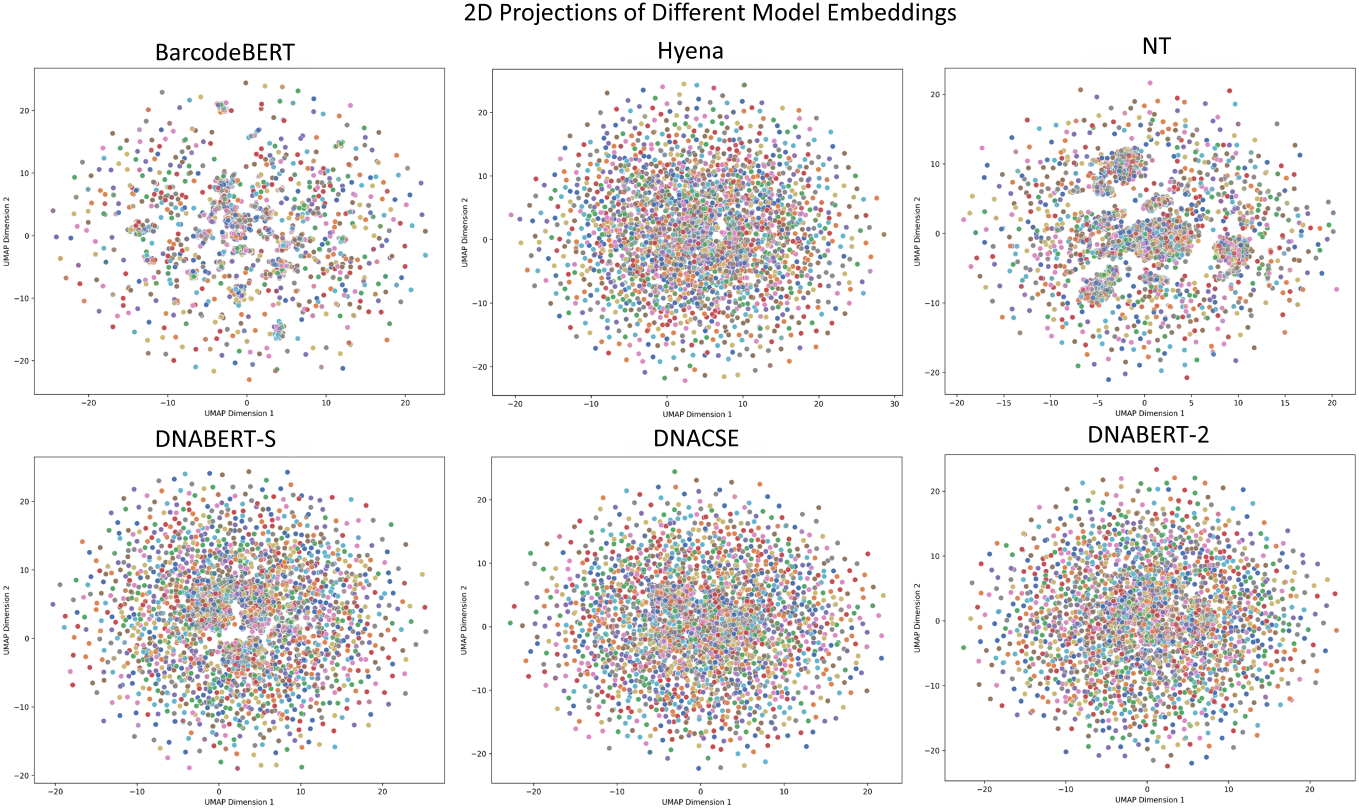
2D projection of different models

### Gene-MTEB Benchmark Tasks

To demonstrate that DNACSE can improve the generalization ability of the model. We evaluated the ability of DNACSE to generate high-quality embeddings in a zero-shot manner with other comparison baselines using the Gene-MTEB^23^benchmark dataset. This benchmark includes eight classification tasks (Human-Virus-1-4, MHPD-single, HMPD-disease, HMPD-source, HMPD-sex) and eight clustering tasks (HVR-p2p, HVR-s2s-align, HVR-s2ssmall, HVR-s2stiny, HMPR-p2p, HMPR-s2s-align, HMPR-s2s-small, HMPR-s2s-tiny). The datasets were obtained from the Human Microbiome Project (HMP),^60^ human viral reference sequences, and genetic sequences of human viral infection samples. Logistic regression was performed using model-generated embeddings for all classification tasks and the small batch k-means algorithm was performed using model-generated embeddings for all clustering tasks. Accuracy was used as the reporting metric for classification tasks, and the v-measure as the evaluation metric for clustering tasks, and we uniformly scaled all metrics by a factor of 100 to represent consistency. Higher scores represent higher performance, which typically means better embedding quality. The mean pooling was used for all models, and only the results of the best-performing model are reported.

Table 2 shows the results of the Gene-MTEB benchmark. Figure 4 shows the average performance of DNACSE and all baseline models in four different types of data: HumanVirus, HMPD, HVR, and HMPR. Due to the unique advantages of DNACSE, the designed feature noise helps the model to generalize to more potential sequence representations. The contrastive learning process enables the model to extract detailed semantic information from the sequences. By further combining the training process with rich pre-trained biological knowledge, higher quality embeddings are generated. We found that DNACSE outperforms the comparison baselines in both the HMPD and HVR tasks, achieving the best overall performance in the four task categories. The final average score improved by 2.8 points compared to the second-best DNABERT-S and by 6.39 points compared to DNABERT-2. These results demonstrate that DNACSE successfully improves the distribution of DNA language model embeddings, thereby enhancing the model’s generalization ability and performance on zero-shot benchmark.

**Table 2:** Model performance on the Gene-MTEB benchmark^*a*^.

| Model/Task | DNABERT-2 | DNABERT-S | NT | Hyena | BarcodeBERT | DNACSE |
| --- | --- | --- | --- | --- | --- | --- |
| Human-Virus-1 | 66.67 | **74.26** | 59.43 | 59.77 | 63.21 | <u>69.69</u> |
| Human-Virus-2 | 49.98 | **61.97** | 57.30 | 51.36 | 61.33 | <u>61.47</u> |
| Human-Virus-3 | 55.04 | <u>67.09</u> | 64.26 | 51.37 | 61.35 | **74.72** |
| Human-Virus-4 | 67.99 | <u>69.28</u> | **69.41** | 65.34 | 61.31 | 66.60 |
| HMPD-single | 28.98 | <u>29.18</u> | 28.75 | 28.27 | 27.25 | **30.27** |
| HMPD-disease | 32.74 | 42.92 | 43.71 | 30.03 | <u>51.69</u> | **60.39** |
| HMPD-sex | 34.64 | 37.10 | 39.16 | 35.29 | <u>48.86</u> | **49.16** |
| HMPD-source | 37.47 | 45.05 | 37.39 | 26.58 | <u>51.72</u> | **66.42** |
| HVR-p2p | <u>59.39</u> | 58.74 | 59.14 | 57.10 | 49.29 | **61.73** |
| HVR-s2s-align | 26.36 | 25.13 | 19.89 | 26.78 | <u>28.13</u> | **28.36** |
| HVR-s2s-small | 37.97 | 33.80 | 39.38 | <u>39.39</u> | 38.30 | **40.31** |
| HVR-s2s-tiny | <u>75.30</u> | <u>75.30</u> | **75.31** | 72.39 | <u>75.30</u> | 70.98 |
| HMPR-p2p | 52.38 | **60.76** | <u>55.05</u> | 53.43 | 53.97 | 53.32 |
| HMPR-s2s-align | 13.37 | **19.00** | 11.44 | 13.09 | 17.28 | <u>18.45</u> |
| HMPR-s2s-small | 41.21 | 40.22 | 37.69 | 39.85 | <u>41.96</u> | **42.99** |
| HMPR-s2s-tiny | **100.00** | 97.08 | 95.63 | **100.00** | <u>98.54</u> | 86.91 |
| Global average | 48.71 | <u>52.30</u> | 49.55 | 46.87 | 51.84 | **55.10** |

<sup>a</sup> Clustering tasks evaluated using v-value, classification tasks using accuracy (both scaled by 100). Best results in **bold**, second-best underlined.

**Figure 4.**
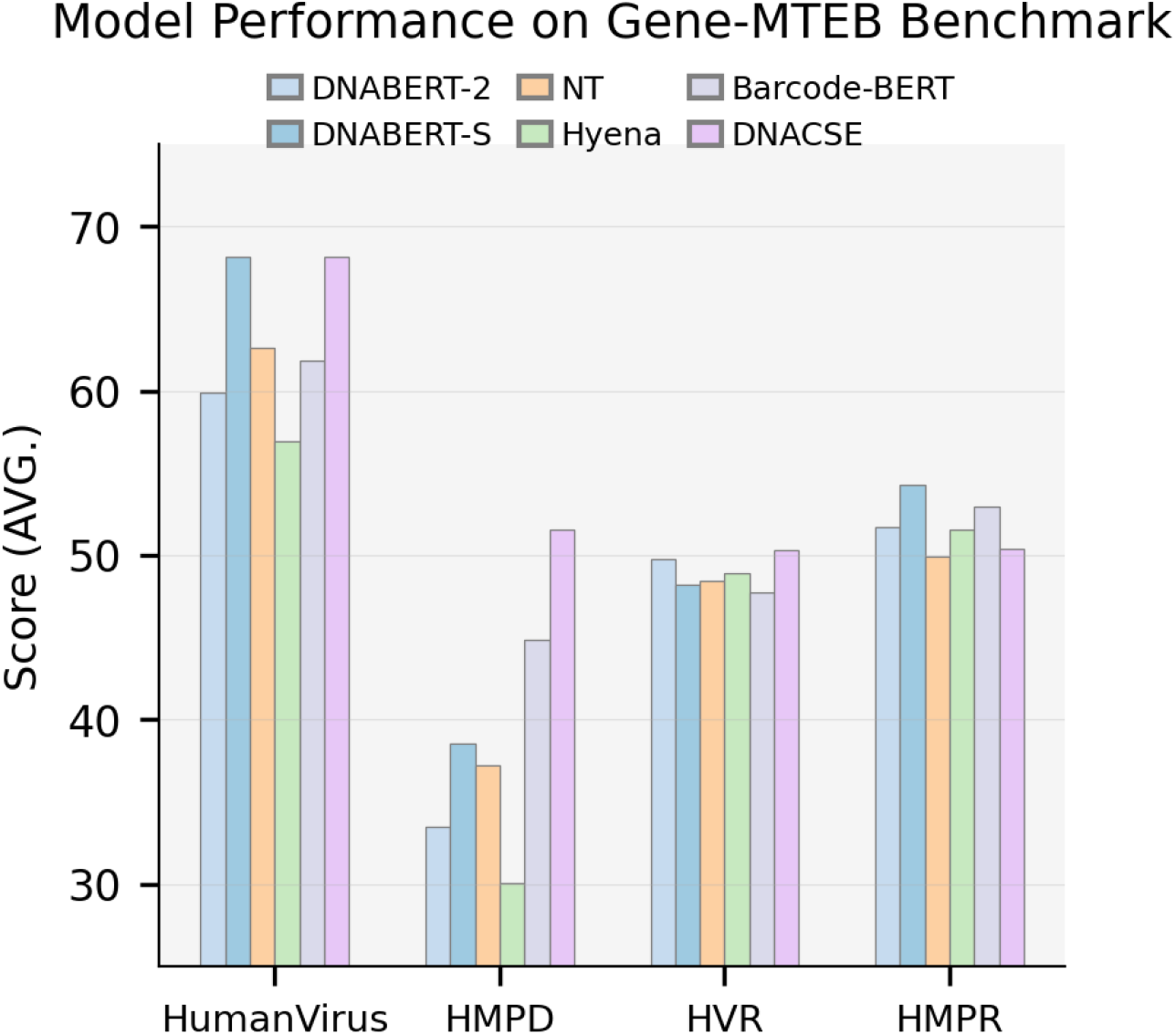
Model performance comparison on Gene-MTEB benchmark across four tasks.

### Genomic Benchmark Tasks

To demonstrate that DNACSE can improve performance in genomic downstream tasks, this paper used Genomic Benchmark,^24^ a combination of new datasets constructed from publicly available database mining and existing datasets obtained from published articles. It contains nine sequence classification tasks consisting of promoters, enhancers, and open chromatin regions of human, mouse, and roundworm, which help to systematically assess the generalization ability of DNA language models in genomic downstream tasks. All of the above tasks use accuracy as a reporting metric. All models use the unified GFMBenchmark^61^ tool to perform 10 rounds of fine-tuning, followed by testing on a test set, and we report the best results from three experiments for all models.

Figure 5 shows the results of the Genomic Benchmark. The grouped chart on the left shows the performance of all models on each task, while the box plot on the right shows the overall distribution and stability of the cross-task performance of all models on the genomic benchmark, higher box lines represent higher accuracy levels, while lower box bodies represent narrower performance fluctuations. Using the noise contrastive learning framework, this approach continuously extracts nuanced hidden features from sequences by pulling positive sample representations closer while pushing away both ordinary and hard negative samples. DNACSE achieved the best level in 5 of all 9 tests, although DNACSE may not be the best performer in all tasks, its overall performance curve is more balanced, indicating that it exhibits stable high performance in most tasks compared to all of the compared models. The results show that DNACSE can effectively improve DNA language modeling in downstream genomic fine-tuning tasks.

**Figure 5.**
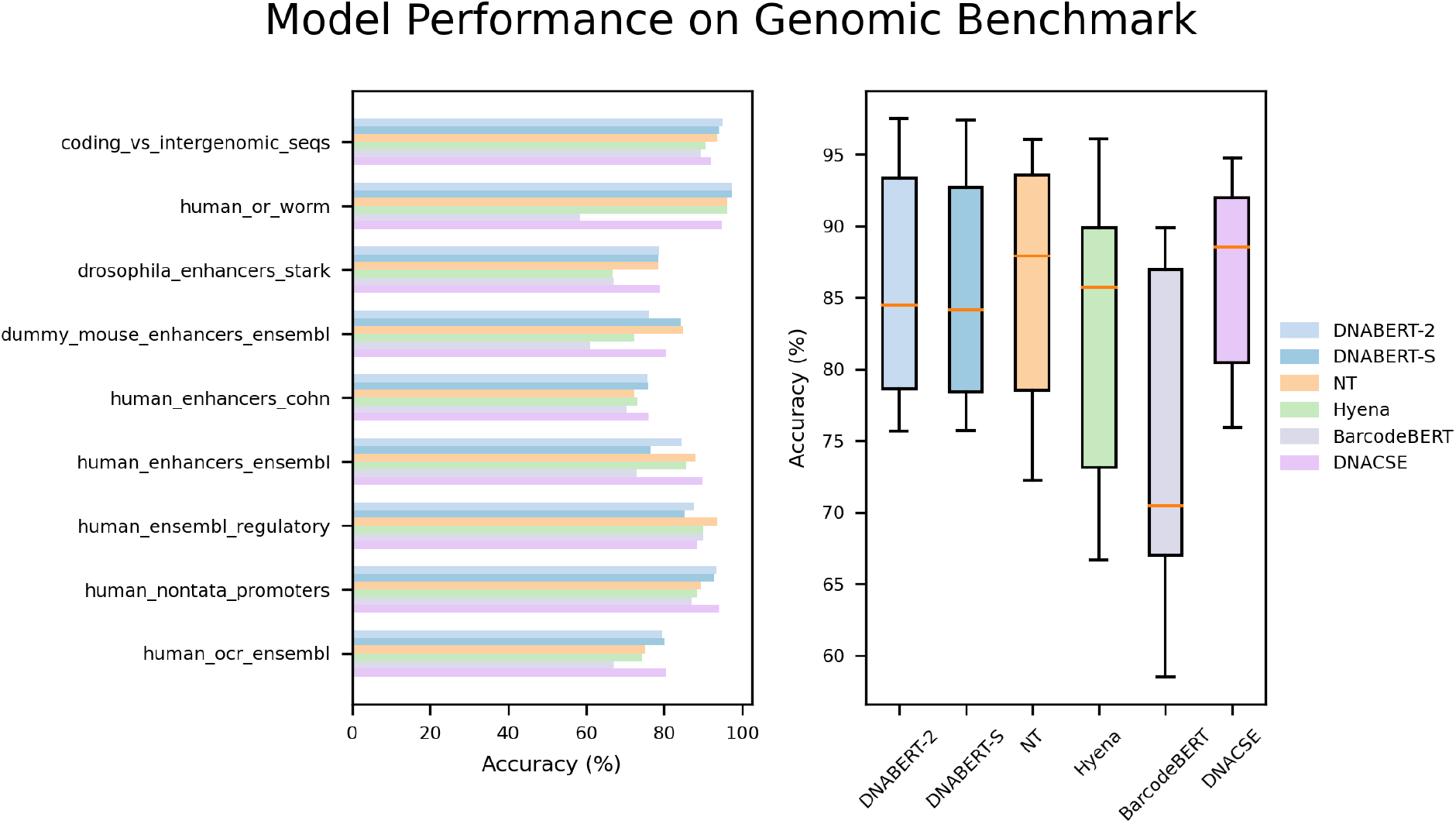
performance of DNACSE and baseline models on all nine different classification tasks in the Genomic Benchmark

### Comparison of Different Data Augmentation Methods

To investigate the most suitable data augmentation method for DNA sequences, we compared several widely used data augmentation methods, including replacing a portion of base positions with the unknown base *N* with a fixed probability of 20% to simulate implicit errors present in sequencing (mask) and performing global rolling shifts on DNA barcodes in non-primer regions to construct different positive sample views (roll).^62^ Furthermore, we tested several feature augmentation methods to replace the original feature noise layer, including randomly discarding (drop) 10% of the feature values in each dimension of the embedding tokens with a fixed probability, using only global noise or using only local noise. We also tested two parallel feature augmentation frameworks:^63^ parallel gaussian and parallel feature dropout. For an anchor sample *z*_*i*_ and its feature-augmented version 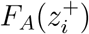, the augmented contrastive loss is calculated as:

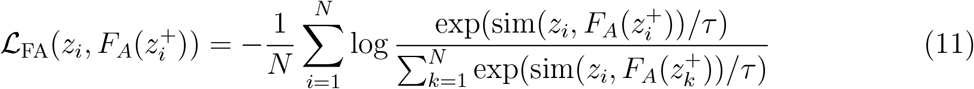

The original contrastive loss is calculated as

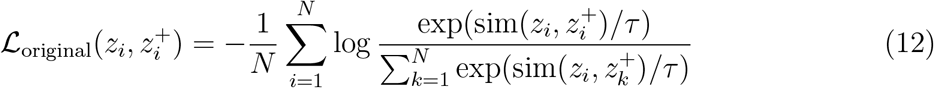

Finally, we average these losses as follows:

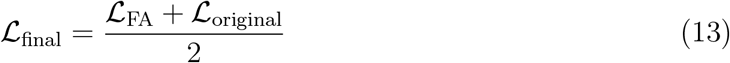

to combine the original view and the feature-augmented view. In this experiment, we used the performance of the BIN reconstruction task as an evaluation metric to assess the performance of various data augmentation methods.

The results in Figure 6 show that the use of feature augmentation as a positive sample pair construction method all outperforms augmentation using sequence-level structural variation. Interestingly, this paper observed that using the *mask&roll* augmentation method yielded significantly better results than using *mask* alone. We hypothesize that the *mask* operation may be too simplistic and, more importantly, risks corrupting key sequence motifs. By only requiring the model to reconstruct masked tokens, this straightforward task might not be challenging enough to compel the model to learn a highly generalized and robust representation. In contrast, the *roll* operation changes the local order of the sequence by cyclically shifting tokens. This modification forces the model to focus not just on individual tokens but also to comprehend the relative positions and global structure between tokens. To correctly identify the original and rolled sequences as positive pairs, the model must learn a representation that is insensitive to such local structural perturbations. This task is inherently more challenging than the simple *mask* operation. Therefore, *mask&roll* combines the advantages of both augmentations: it introduces local semantic disruption via masking while simultaneously adding a global structural challenge via rolling. This combination creates a more balanced and demanding training objective, which in turn prompts the model to learn more discriminative and robust feature representations, ultimately leading to a significant performance boost. In contrast, We argue that feature-level augmentation, with more nuanced perturbations, carries less risk of corrupting the semantics when constructing positive sample pairs compared to sequence-level. In addition, the combined use of feature noise outperforms the performance achieved by either method alone. This might be because the feature augmentation approach perturbs both the direction and magnitude of features, creating a challenging yet manageable training target. Therefore, this process results in improved embeddings, further demonstrating the effectiveness of the method.

**Figure 6.**
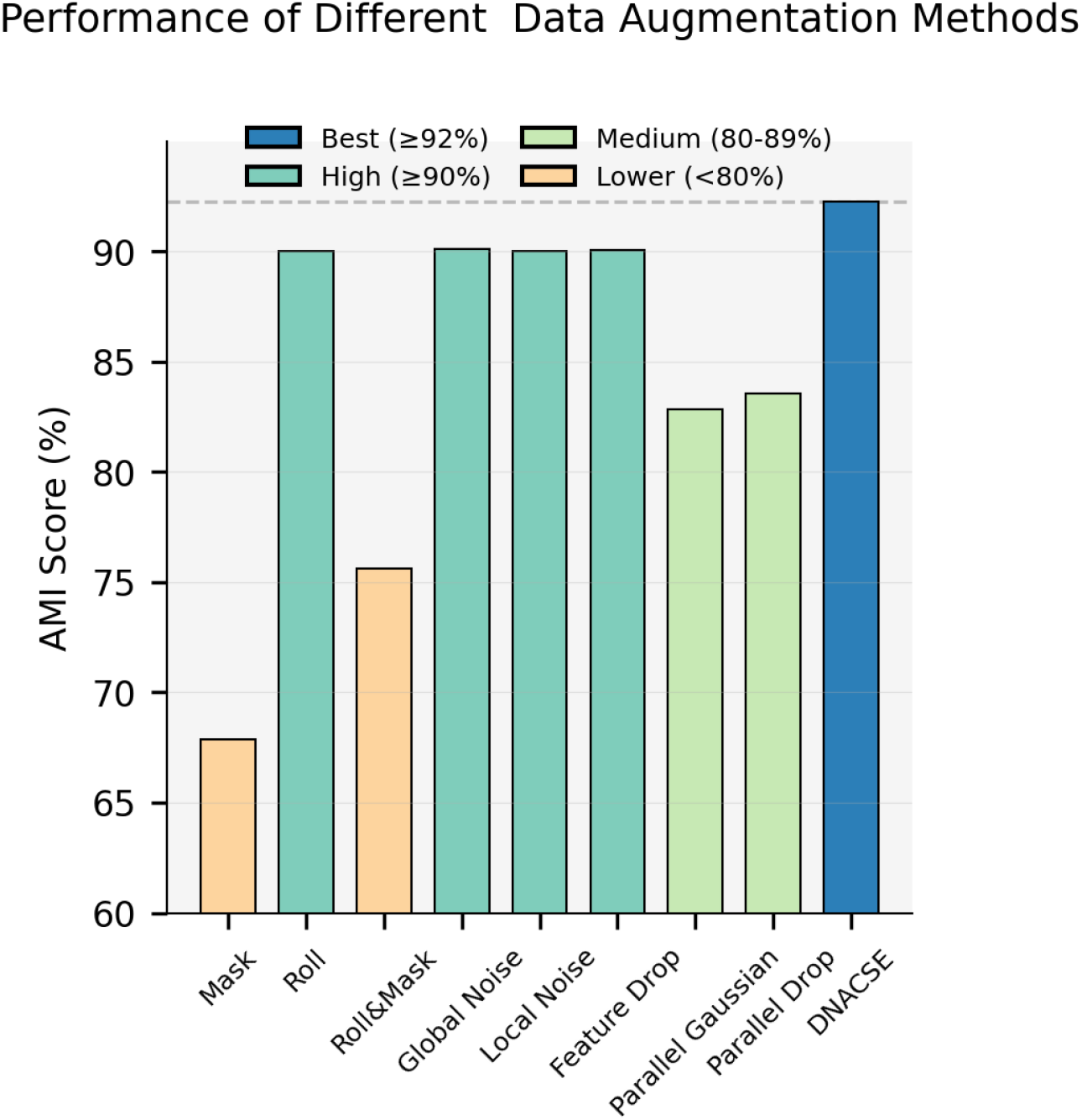
Performance comparison of different data augmentation methods on AMI metric for BIN reconstruction task.

### DNACSE Enhances Representation Space Isotropy and Separability

To systematically evaluate and compare the performance of different models in DNA barcode embedding, this paper employed a series of representation learning metrics, ^59^ including alignment, uniformity, and separability. These metrics provide an intuitive and effective reflection of the models’ characteristics in terms of distribution within the embedding space. The experimental results are shown in Figure 7. DNACSE significantly outperformed all baseline models, including its foundational model DNABERT-2, across key metrics, fully demonstrating its effectiveness in optimizing the quality of DNA representation spaces. As observed in the scatter plot of Figure 7A, DNACSE successfully avoids the issues of over-alignment and over-uniformity faced by other models. Models like NT, DNABERT-2, and DNABERT-S tend to over-align, leading to excessive clustering of class embeddings and thereby weakening their ability to distinguish between classes. BarcodeBERT, however, goes to the opposite extreme by excessively pursuing uniformity. This forces embeddings to scatter across the hypersphere, flattening natural inter-class differences and hindering effective clustering of samples within the same class. Different from these models, DNACSE achieves superior uniformity while maintaining good alignment. DNACSE achieves a uniformity score of -2.5657, significantly outperforming DNABERT-2, DNABERT-S, and other baseline models. This indicates that its generated embeddings exhibit more uniform distribution across the unit hypersphere while maintaining strong discriminative power, effectively mitigating anisotropy issues in the representation space.

**Figure 7.**
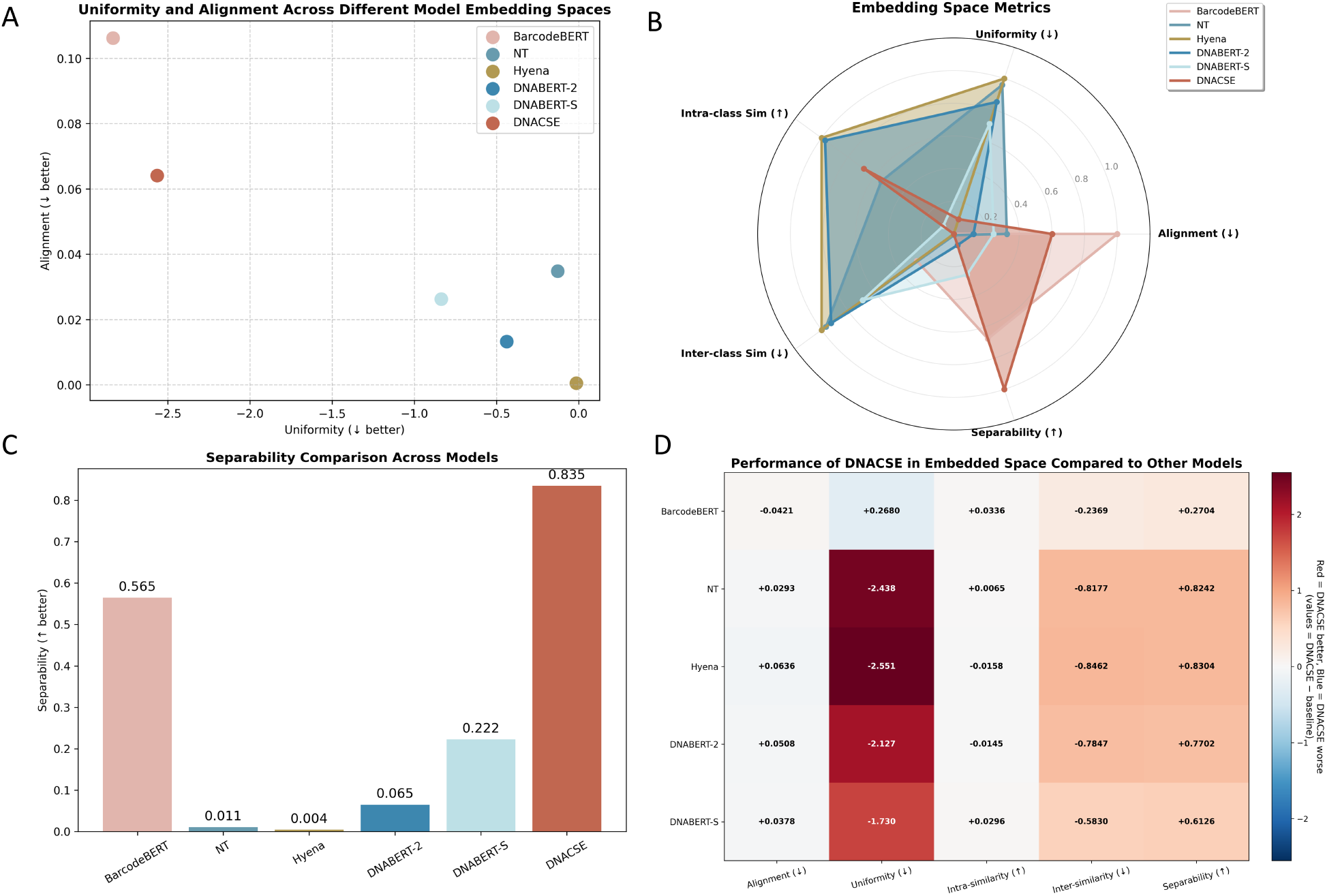
Performance Across Different Model Embedding Spaces. (A) Uniformity and Alignment across different model embedding spaces, with lower values indicating better performance. (B) Radar chart of embedding space metrics including intra-class similarity, inter-class similarity, uniformity, alignment, and separability for various models. (C) Bar chart showing separability across different model embedding spaces; higher values indicate better performance. (D) Heatmap showing the performance gap between DNACSE and other models, covering metrics such as alignment quality, uniformity, internal similarity, external similarity, and separability.

Figure 7B’s radar chart further summarizes the comparison across multiple metrics, revealing that DNACSE demonstrates a prominent advantage in the Separability dimension, reflecting its comprehensive performance in constructing high-quality feature spaces. The bar chart in Figure 7C and the performance variance heatmap in Figure 7D quantify this advantage: Compared to DNABERT-2, DNACSE achieves a 0.7702 improvement in Separability and a 2.127 enhancement in Uniformity, demonstrating significant performance gains.

To gain deeper insights into the characteristics of the embedded space revealed by these metrics, this paper further visualized the cosine similarity matrices and the distributions of inter-class and intra-class similarities for DNACSE, DNABERT-2, and DNABERT-S. The visualized results are shown in Figure 8. The ideal representation space should manifest in the cosine similarity matrix as bright, sharply defined diagonal blocks contrasted against overall dark non-diagonal regions, The heatmap of DNACSE (Figure 8C) effectively illustrates this ideal structure, whereas the heatmaps of DNABERT-S (Figure 8A) and DNABERT-2 (Figure 8B) exhibit an overall reddish bias with insufficient contrast between non-diagonal and diagonal regions. This indicates that the models fail to effectively discriminate samples across different classes, resulting in unreasonably high similarity between all sample pairs—a classic visual manifestation of anisotropy in the representation space. The subsequent similarity distribution histogram further quantifies this observation. The distribution of intra-class similarity for DNACSE clusters around the high-value region near 1.0, while the inter-class similarity distribution clusters in a lower range. A distinct gap exists between the two distributions, demonstrating outstanding separability. In contrast, the intra-class and inter-class similarity distributions for DNABERT-2 overlap severely, making them nearly inseparable. DNABERT-S (Figure 8D) shows a similar but slightly less severe overlap, which directly accounts for its lower separability score.

**Figure 8.**
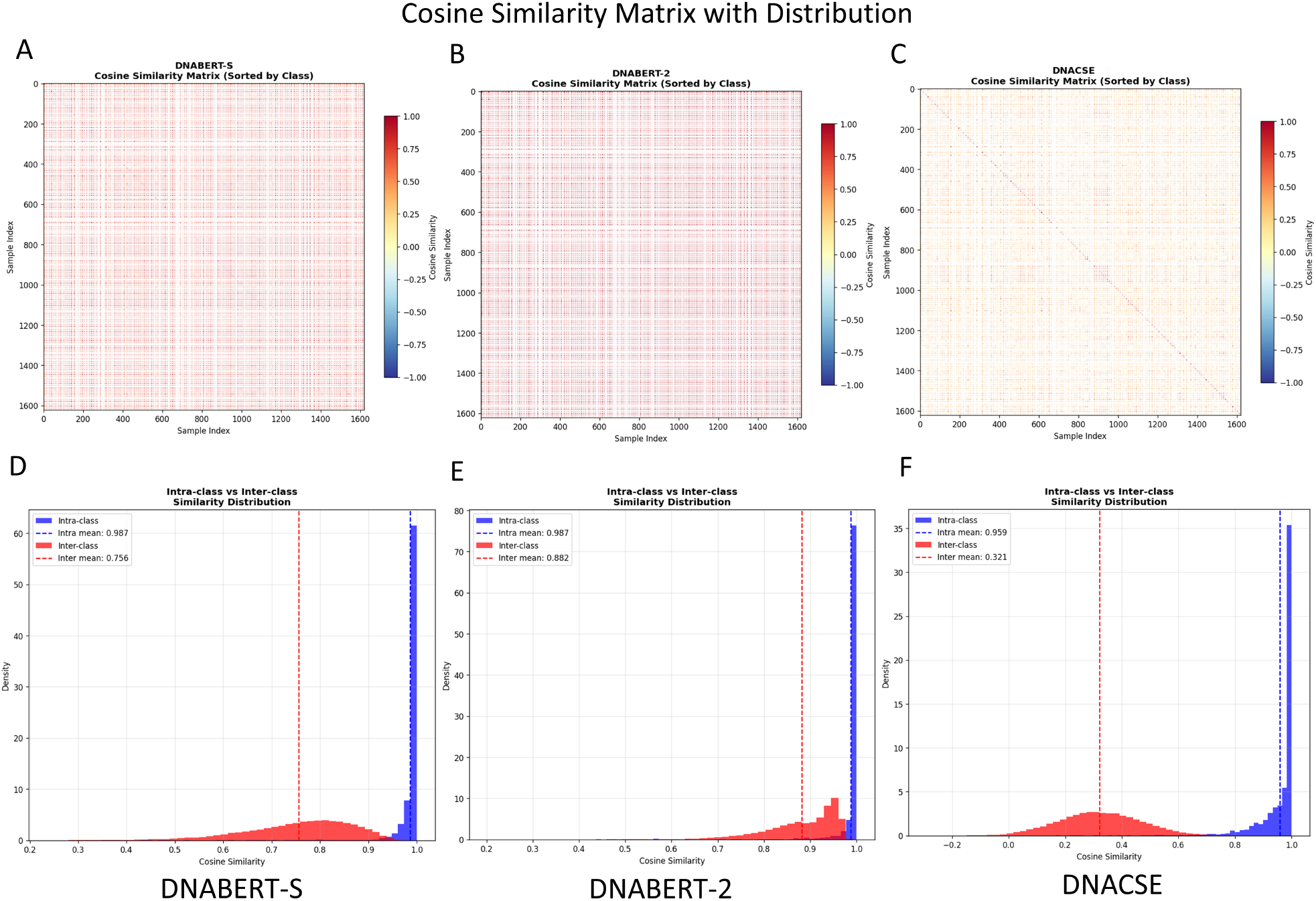
Cosine Similarity Matrix with Distribution Across Different Models. (A) Cosine similarity matrix for DNABERT-S sorted by class. (B) Cosine similarity matrix for DNABERT-2 sorted by class. (C) Cosine similarity matrix for DNACSE sorted by class. (D) Intra-class vs inter-class similarity distribution for DNABERT-S. (E) Intra-class vs interclass similarity distribution for DNABERT-2. (F) Intra-class vs inter-class similarity distribution for DNACSE, with mean values indicating the average similarity within and between classes.

In summary, both the quantitative metrics and the qualitative visualizations of the embedding space consistently and convincingly demonstrate DNACSE’s effectiveness in enhancing representation uniformity and improving class separability.

### Generalizability across Different Models

To further validate the generalization capability of our framework, this paper extended the evaluation beyond DNABERT-2, which was used as the base model in the main experiments. This paper incorporated three additional pre-trained sequence models with distinct architectures and training corpora: NT, BarcodeBERT and Hyena These models vary in design philosophy and training data scale, providing a comprehensive benchmark for assessing the adaptability of DNACSE. To ensure consistency in our results, this paper used a unified noise configuration (*globalstd* = 0.04, *localstd*=0.15, *mutationrate* = 0.1) and consistent training settings, including temperature *τ* =0.07 and margins *m* = 0.02. this paper used the BIN reconstruction task as the evaluation metric. For both NT and BarcodeBERT, this paper employed hybrid pooling strategy for training and used average pooling for evaluation. Since Hyena lacks a CLS token, this paper used average pooling for both training and evaluation.

Figure 9 summarizes the experimental results on the BIN reconstruction task. Across all settings, DNACSE consistently improves the baseline performance of each model, regardless of architectural differences. These results strongly support the claim that DNACSE is broadly applicable and not limited to DNABERT-2. By validating its effectiveness across diverse base encoders, we establish the outstanding generalizability of DNACSE.

**Figure 9.**
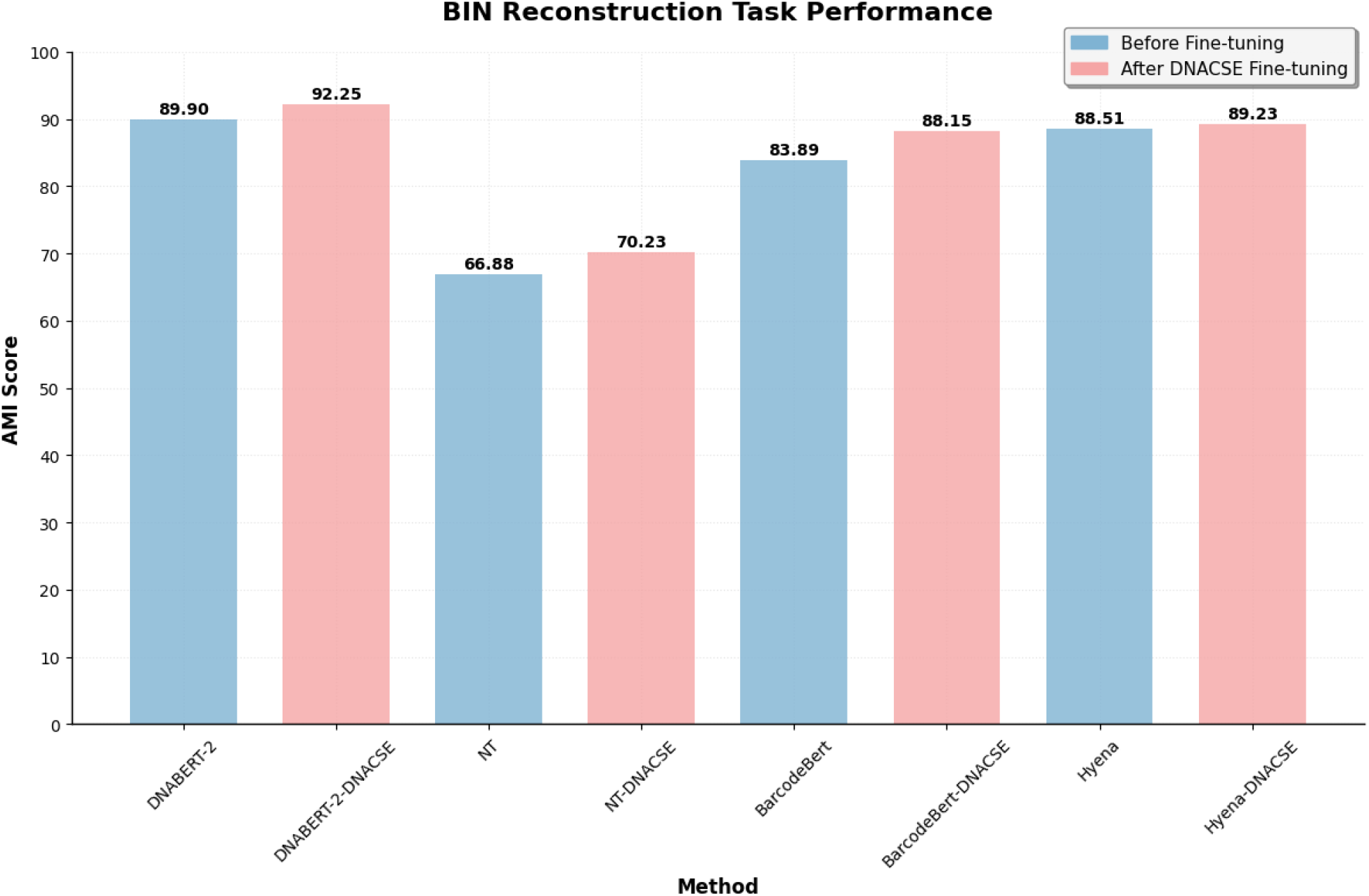
AMI Score Comparison Before and After DNACSE Fine-tuning

### Ablation Study

In order to demonstrate the effectiveness of each component of DNACSE. This paper chooses BIN reconstruction as the evaluation criterion and performed a thorough ablation of each DNACSE component. We established a baseline model using standard unsupervised contrastive learning, where positive pairs were constructed solely from the sequence itself. We then sequentially evaluated the contributions of individual components by adding the feature noise module, the hard negative sample mixing module, and finally integrating all modules to form the complete DNACSE model. Generally, all sub-components use average pooling as the final pooling method in ablation experiments.

Table 3 shows the ablation results of each component of DNACSE, and the results indicate that the feature noise module layer and the difficult negative sample mixing module construction strategy can effectively improve the performance on BIN reconstruction tasks, which may be due to the fact that the feature noise module somewhat introduces more detailed perturbations for the positive sample pairs, forcing the model to learn the noise-invariant representations, while the hard negative sample mixing module can force the model to further explore features that are prone to non-recognition, enhancing the model’s performance. These results further demonstrate the reliability of feature data augmentation and difficult negative samples in improving model performance.

**Table 3:**
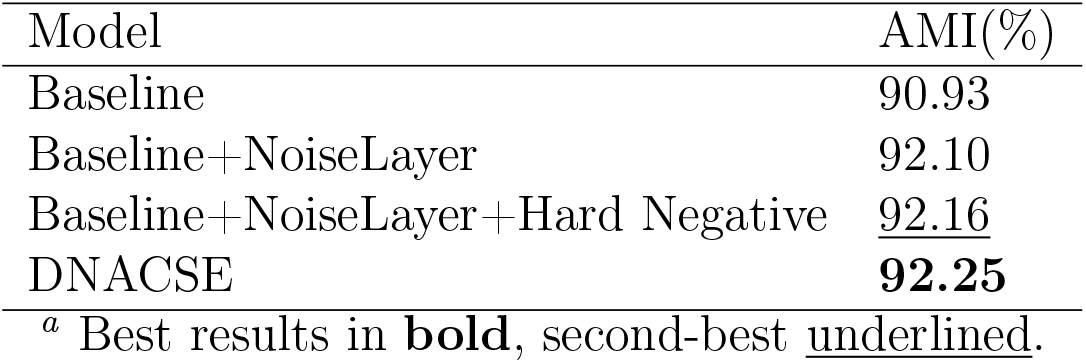
Performance comparison of different model configurations on AMI metric^*a*^.

| Model | AMI(%) |
| --- | --- |
| Baseline | 90.93 |
| Baseline+NoiseLayer | 92.10 |
| Baseline+NoiseLayer+Hard Negative | <u>92.16</u> |
| DNACSE | **92.25** |
| <sup>a</sup> Best results in **bold**, second-best <u>underlined</u>. |  |

To explore the influence of different batch sizes and pooling methods on DNACSE’s performance, we tested several common pooling methods, including the cls token, average pooling (avg) and the average of the first and last layer embeddings (first_last_avg). Additionally, we implemented a hybrid pooling method (mix) that alternates between cls and average pooling during training. For the cls token, the original BERT implementation used an additional MLP layer. Here, we considered two implementations of cls: 1) We removed this MLP layer and used the original cls token as the pooling method (cls before pooler). 2) Maintain the MLP layer (cls). Table 4 shows the performance of DNACSE in the BIN reconstruction task under different pooling methods.

**Table 4:**
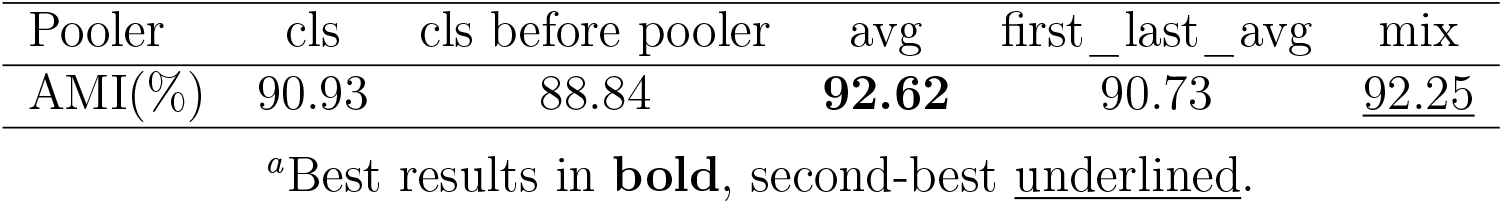
Performance of different pooling methods.

| Pooler | cls | cls before pooler | avg | first_last_avg | mix |
| --- | --- | --- | --- | --- | --- |
| AMI(%) | 90.93 | 88.84 | **92.62** | 90.73 | <u>92.25</u> |

“Best results in **bold**, second-best underlined.”

Compared to other pooling methods, average pooling aggregates the structural information of each token on average, enabling it to better capture global information and reflect the distribution characteristics of the input sequence. This is consistent with the objective of clustering the task of grouping based on feature similarity. Interestingly, although we found that average pooling is more suitable for clustering tasks, models trained with mix pooling achieved better performance than trained with average pooling in classification on other tasks. We believe that the mix pooling method helps the model capture global information while enhancing its ability to encode global semantics to some extent. Specifically, Our mix pooling method does not simply average features; it integrates the complementary strengths of global semantic information and local salient Information. Therefore, mix pooling creates a richer, more comprehensive representation that captures both the global semantic context and the key local details. This dual-encoding capability provides the model with a more robust and versatile set of features. While a global average is sufficient for clustering based on broad similarities, a combination of global and salient features allows the model to better handle the nuances and discriminative features required by other tasks, such as classification, where the model needs to distinguish between different categories with high precision. This enhanced representational power is what enables the improved generalization ability observed in our experiments.

As shown in Table 5, the influence of different batch sizes on the BIN reconstruction task. Since Table 4 illustrates that the average pooling is the best performing pooling method for the BIN reconstruction task, we only use the average pooling as the evaluation pooling method. With increasing batch size, the performance of the BIN reconstruction task improves slightly, but the overall trend remains stable, and memory usage also increases significantly. When the batch size exceeds 128, this improvement begins to decline. Therefore, we believe that 128 achieves the optimal balance between performance and computational burden in this scenario. When the batch size varies within the tested range, the model’s clustering performance demonstrates high robustness. This indicates that in DNACSE, model performance is primarily determined by the data augmentation strategy and the model’s inherent stability, rather than simply depending on the batch size.

**Table 5:** Performance of different batch sizes.

| Batch size | 32 | 64 | 128 | 256 |
| --- | --- | --- | --- | --- |
| AMI(%) | <u>92.56</u> | 92.46 | **92.62** | 92.50 |

<sup>a</sup>Best results in **bold**, second-best underlined.

In order to explore the impact of different temperatures and margins on the training dynamics of DNACSE, we conducted a series of ablation studies by testing various combinations of these two parameters. Following the principle of a controlled experiment, this paper fixed one parameter at a value that demonstrated reliable performance in preliminary experiments and then systematically examined the independent effect of the other. Specifically, this paper fixed *m* = 0.2 when examining *τ*, and this paper fixed *τ* = 0.08 when examining *m*. To ensure the comparability of our results across all experiments, this paper maintained consistent settings for noise hyperparameters (*globalstd* = 0.04, *localstd* = 0.15, *mutationrate* = 0.10), training configurations, CLS pooling methods, and the evaluation metrics for the BIN reconstruction task.

Table 6 details the impact of different temperatures on the performance of DNACSE in the BIN reconstruction task under fixed marginal values, where Steps records the number of evaluation steps required to achieve optimal performance. Different temperatures *τ* reflect the impact of temperature on convergence rate and peak performance. At lower temperatures, the distribution becomes overly sharp, with negative samples exhibiting excessive discriminative power that disrupts positive-negative balance, resulting in lower AMI. Conversely, moderate temperatures (e.g., *τ* =0.07) strike a balance between discriminative power and balance, achieving optimal performance within fewer training steps. larger temperatures (e.g., *τ* =0.08) smooth the distribution, resulting in slower improvement but ultimately yielding good results. This indicates that *τ* primarily regulates convergence speed and the separation degree of the embedding space.

**Table 6:** Performance of Different Temperatures in the BIN Reconstruction Task^*a*^.

| $\tau$ | 0.05 | 0.06 | 0.07 | 0.08 |
| --- | --- | --- | --- | --- |
| AMI (%) | 88.43 | 88.82 | **90.93** | <u>90.26</u> |
| Steps | 3250 | 5000 | 12000 | 12250 |

<sup>a</sup>Best results in **bold**, second-best underlined.

Table 7 details the impact of different threshold values on the performance of DNACSE in the BIN reconstruction task under fixed temperature conditions. Different margin values *m* reflect the regulatory effect of margin on final performance and convergence stability. Smaller m (e.g., *m*=0.01) enables rapid learning but results in a loosely structured embedding space, limiting the upper bound of performance; Moderate *m* (e.g., *m*=0.03) converges to peak performance within more steps (around 12500) and maintains stability; whereas larger *m* excessively compresses positive sample distances, slowing learning and degrading performance. This indicates *m* primarily influences the internal structure of embeddings and model training stability. Overall, Tables 6 and 7 reveal the complementary effects of temperature and margin on learning performance from different dimensions: temperature determines the separation and convergence speed of the embedding space, while margin influences the aggregation of positive samples and training stability.

**Table 7:** Performance of Different Margins in the BIN Reconstruction Task^*a*^.

| $m$ | 0.01 | 0.02 | 0.03 | 0.04 | 0.05 |
| --- | --- | --- | --- | --- | --- |
| AMI (%) | 85.83 | **90.93** | 90.73 | 90.34 | <u>90.34</u> |
| Steps | 5000 | 12000 | 12500 | 17500 | 12000 |

<sup>a</sup>Best results in **bold**, second-best underlined.

## Discussion and Conclusion

This paper introduces the DNACSE, which high-efficiency fine-tunes DNA language models through unsupervised contrastive learning. The framework combines feature noise layers with hard negative sample construction strategies to better understand DNA barcodes patterns. This paper aims to leverage rich multi-species pre-trained parameters as background knowledge to fully unlock the model’s potential on genomic biodiversity data, particularly in DNA barcoding species taxonomic tasks. We evaluated the performance of DNACSE across multiple DNA barcoding tasks and compared it with several language models pretrained on non-DNA or DNA barcoding data. In addition, we compared the quality of the embeddings generated by each model and their transferability to zero-shot and fine-tuning benchmarks. Finally, we explored the most suitable data augmentation methods for DNA barcoding contrastive learning through a series of ablation experiments and analyzed the influence of different configurations of DNACSE.

In DNA barcoding tasks, DNACSE outperformed other DNA language models and almost achieved the best results. In particular, DNACSE made significant progress in linear probe and zero-shot clustering tasks, indicating that it can generate unique and high-quality DNA barcode embeddings. Additionally, DNACSE has demonstrated its superiority in finetuning tasks by addressing domain shift issues by learning the deep semantics of DNA barcodes, whereas traditional supervised fine-tuning models struggle to achieve the same level of performance. These remarkable results are primarily attributed to two key factors: 1) An optimized unsupervised contrastive learning objective. By pulling semantically similar sequences closer together and pushing unrelated sequences apart, the model can effectively cluster closely related species in the feature space.^64,65^ Feature noise helps the model generalize, while mixed hard negative samples assist the model in identifying differences between real and pseudo DNA barcodes, thereby enhancing its ability to distinguish subtle differences between genuine and pseudo DNA barcodes. 2) Rich pre-training background information. DNACSE leverages rich multi-species information as implicit guidance, enabling it to gain a deeper understanding of DNA barcode data patterns, outperforming models pre-trained solely on DNA barcode data.^66,67^ DNA barcode information provides DNACSE with a clear optimization direction, enabling it to demonstrate stronger adaptability to such data, outperforming models trained on non-DNA barcode data.

The results of zero-shot and fine-tuning benchmarks prove that DNACSE helps improve the distribution of DNA language model embeddings, thereby producing high-quality embeddings. Furthermore, DNACSE enhances the model’s transferability, leading to improved performance in both zero-shot and fine-tuning tasks. Although DNACSE does not always perform optimally in the fine-tuning benchmark, the box plot in Figure 5 shows that DNACSE exhibits smaller performance gaps between tasks compared to other models, suggesting its understated robustness. We believe that this performance is the result of the DNACSE training process adjusting model parameters to better suit DNA barcode data, thus validating the effectiveness of the training to some extent. We also compared various explicit and feature-based data augmentation methods, and the results showed that feature augmentation achieved the best results. This not only demonstrates its effectiveness, but also indicates that, in biological sequence contrastive learning, feature-level data augmentation is simpler, carries lower risk, and yields more significant optimization effects compared to higher-risk explicit augmentation techniques.^68^ This finding provides new insights for the exploration of DNA barcoding. Finally, through exhaustive ablation experiments, ablation studies confirmed that the introduction of the feature noise layer and the hard negative sample construction strategy improved the model’s performance in zero-shot clustering. Future work will focus on extending the DNA language model to more generalized and universal datasets and investigating how to mitigate the representation degradation issues of DNA language models.

## Data and Software Availability statement

As described above, https://github.com/Kavicy/DNACSE is open-source and freely available on the GitHub repository. The data set used to test the functionality of the tool and additional installation information is available in the same repository.

## Author Contributions

Conceptualization, Jiadong Wang, Ben Cao, Bin Wang and Shihua Zhou; methodology, Jiadong Wang, Ben Cao; investigation, Wei Li; writing—original draft, Jiadong Wang and Ben Cao; writing—review & editing, Bin Wang and Pan Zheng; supervision, Bin Wang, Shihua Zhou, Ben Cao; funding acquisition, Bin Wang and Shihua Zhou.

## Acknowledgement

The authors thank the Key Laboratory of Advanced Design and Intelligent Computing for providing the necessary facilities and equipment to carry out our experiments. The authors also appreciate the guidance and assistance provided by the lab technicians and staff members throughout the experimental process.

## TOC Graphic

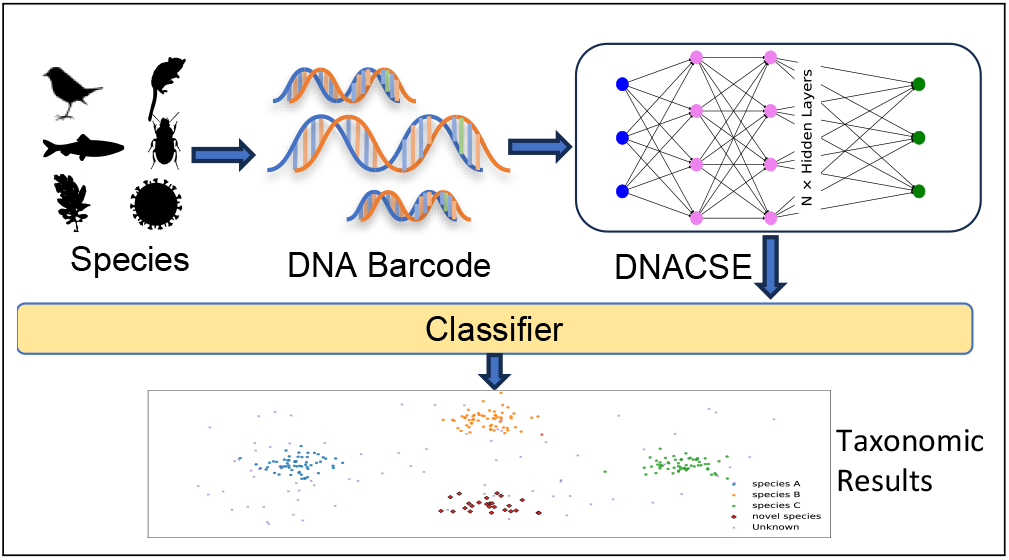

## References

(1) Kress, W. J.; Erickson, D. L. DNA barcodes: genes, genomics, and bioinformatics. Proceedings of the National Academy of Sciences 2008, 105, 2761–2762.

(2) Ruppert, K. M.; Kline, R. J.; Rahman, M. S. Past, present, and future perspectives of environmental DNA (eDNA) metabarcoding: A systematic review in methods, monitoring, and applications of global eDNA. Global Ecology and Conservation 2019, 17, e00547.

(3) Stoeck, T.; Frühe, L.; Forster, D.; Cordier, T.; Martins, C. I.; Pawlowski, J. Environmental DNA metabarcoding of benthic bacterial communities indicates the benthic footprint of salmon aquaculture. Marine Pollution Bulletin 2018, 127, 139–149.

(4) Barcaccia, G.; Lucchin, M.; Cassandro, M. DNA barcoding as a molecular tool to track down mislabeling and food piracy. Diversity 2015, 8, 2.

(5) Gizaw, Z. Public health risks related to food safety issues in the food market: a systematic literature review. Environmental health and preventive medicine 2019, 24, 68.

(6) Hebert, P. D.; Cywinska, A.; Ball, S. L.; DeWaard, J. R. Biological identifications through DNA barcodes. Proceedings of the Royal Society of London. Series B: Biological Sciences 2003, 270, 313–321.

(7) Schoch, C. L.; Seifert, K. A.; Huhndorf, S.; Robert, V.; Spouge, J. L.; Levesque, C. A.; Chen, W.; Consortium, F. B.; List, F. B. C. A.; Bolchacova, E.; others Nuclear ribosomal internal transcribed spacer (ITS) region as a universal DNA barcode marker for Fungi. Proceedings of the national academy of Sciences 2012, 109, 6241–6246.

(8) Letsiou, S.; Madesis, P.; Vasdekis, E.; Montemurro, C.; Grigoriou, M. E.; Skavdis, G.; Moussis, V.; Koutelidakis, A. E.; Tzakos, A. G. DNA barcoding as a plant identification method. Applied Sciences 2024, 14, 1415.

(9) Altschul, S. F.; Gish, W.; Miller, W.; Myers, E. W.; Lipman, D. J. Basic local alignment search tool. Journal of molecular biology 1990, 215, 403–410.

(10) Nugent, C. M.; Adamowicz, S. J. Alignment-free classification of COI DNA barcode data with the Python package Alfie. Metabarcoding and Metagenomics 2020, 4, e55815.

(11) Das, R.; Rai, A.; Mishra, D. C. CNN_FunBar: advanced learning technique for fungi ITS region classification. Genes 2023, 14, 634.

(12) Jin, L.; Yu, J.; Yuan, X.; Du, X. Fish classification using DNA barcode sequences through deep learning method. Symmetry 2021, 13, 1599.

(13) Wu, J.; Wang, P.; Zheng, Y.; Wang, B.; Zhang, Q.; Zheng, P. Stable DNA Storage Encoding Scheme Based on Repeating Substring Tree. IEEE Transactions on Computational Biology and Bioinformatics 2025,

(14) Sun, C.; Zhang, L.; Zhang, L.; Song, Y.; Ma, B.; Wang, Y. GATRsite: RNA–Ligand Binding Site Prediction Using Graph Attention Networks and Pretrained RNA Language Models. Journal of Chemical Information and Modeling 0, 0, null, PMID: 40814145.

(15) Wang, S.; Bergman, M. T.; Hall, C. K.; You, F. De Novo Design of Multiple Microplastic-Binding Peptides with a Protein Language Model-Guided Generative Adversarial Network. Journal of Chemical Information and Modeling 0, 0, null, PMID: 40765481.

(16) Arias, P. M.; Sadjadi, N.; Safari, M.; Gong, Z.; Wang, A. T.; Lowe, S. C.; Haurum, J. B.; Zarubiieva, I.; Steinke, D.; Kari, L.; others BarcodeBERT: Transformers for biodiversity analysis. arXiv preprint 2311.02401 2023,

(17) Safari, M.; Arias, P. M.; Lowe, S. C.; Kari, L.; Chang, A. X.; Taylor, G. W. Enhancing DNA Foundation Models to Address Masking Inefficiencies. arXiv preprint 2502.18405 2025,

(18) Gao, T.; Taylor, G. W. BarcodeMamba: State space models for biodiversity analysis. arXiv preprint 2412.11084 2024,

(19) Gao, J.; He, D.; Tan, X.; Qin, T.; Wang, L.; Liu, T.-Y. Representation degeneration problem in training natural language generation models. arXiv preprint 1907.12009 2019,

(20) Kalantidis, Y.; Sariyildiz, M. B.; Pion, N.; Weinzaepfel, P.; Larlus, D. Hard negative mixing for contrastive learning. Advances in neural information processing systems 2020, 33, 21798–21809.

(21) Zhang, Y.; Zhang, R.; Mensah, S.; Liu, X.; Mao, Y. Unsupervised sentence representation via contrastive learning with mixing negatives. Proceedings of the AAAI Conference on Artificial Intelligence. 2022; pp 11730–11738.

(22) Vinh, N. X.; Epps, J.; Bailey, J. Information theoretic measures for clusterings comparison: is a correction for chance necessary? Proceedings of the 26th annual international conference on machine learning. 2009; pp 1073–1080.

(23) Liu, O.; Jaghouar, S.; Hagemann, J.; Wang, S.; Wiemels, J.; Kaufman, J.; Neiswanger, W. Metagene-1: Metagenomic foundation model for pandemic monitoring. arXiv preprint 2501.02045 2025,

(24) Grešová, K.; Martinek, V.; Čechák, D.; Šimeček, P.; Alexiou, P. Genomic benchmarks: a collection of datasets for genomic sequence classification. BMC Genomic Data 2023, 24, 25.

(25) Gao, T.; Yao, X.; Chen, D. Simcse: Simple contrastive learning of sentence embeddings. arXiv preprint 2104.08821 2021,

(26) Zhou, Z.; Ji, Y.; Li, W.; Dutta, P.; Davuluri, R.; Liu, H. Dnabert-2: Efficient foundation model and benchmark for multi-species genome. arXiv preprint 2306.15006 2023,

(27) Hendrycks, D.; Gimpel, K. Gaussian error linear units (gelus). arXiv preprint 1606.08415 2016,

(28) Chen, T.; Kornblith, S.; Norouzi, M.; Hinton, G. A simple framework for contrastive learning of visual representations. International conference on machine learning. 2020; pp 1597–1607.

(29) Gharaee, Z.; Lowe, S. C.; Gong, Z.; Millan Arias, P.; Pellegrino, N.; Wang, A. T.; Haurum, J. B.; Eyriay, I.; Kari, L.; Steinke, D.; others BIOSCAN-5M: a multimodal dataset for insect biodiversity. Advances in Neural Information Processing Systems 2024, 37, 36285–36313.

(30) Chen, B.; Sultan, M. M.; Karaletsos, T. Compositional deep probabilistic models of dna-encoded libraries. Journal of Chemical Information and Modeling 2024, 64, 1123– 1133.

(31) Wang, K.; Cao, B.; Ma, T.; Zhao, Y.; Zheng, Y.; Wang, B.; Zhou, S.; Zhang, Q. Storing Images in DNA via base128 Encoding. Journal of Chemical Information and Modeling 2024, 64, 1719–1729.

(32) Liu, Z.; Cao, B.; Shao, Q.; Zheng, Y.; Wang, B.; Zhou, S.; Zheng, P. Family of Mutually Uncorrelated Codes for DNA Storage Address Design. IEEE Transactions on NanoBioscience 2025,

(33) Zhou, Z.; Wu, W.; Ho, H.; Wang, J.; Shi, L.; Davuluri, R. V.; Wang, Z.; Liu, H. DNABERT-S: Pioneering species differentiation with species-aware DNA embeddings. Bioinformatics 2025, 41, i255–i264.

(34) Press, O.; Smith, N. A.; Lewis, M. Train short, test long: Attention with linear biases enables input length extrapolation. arXiv preprint 2108.12409 2021,

(35) Wu, C.; Wu, F.; Huang, Y. Rethinking infonce: How many negative samples do you need? arXiv preprint 2105.13003 2021,

(36) Wu, J.; Chen, J.; Wu, J.; Shi, W.; Wang, X.; He, X. Understanding contrastive learning via distributionally robust optimization. Advances in Neural Information Processing Systems 2023, 36, 23297–23320.

(37) Meyer, C. P.; Paulay, G. DNA barcoding: error rates based on comprehensive sampling. PLoS biology 2005, 3, e422.

(38) Wiemers, M.; Fiedler, K. Does the DNA barcoding gap exist?–a case study in blue butterflies (Lepidoptera: Lycaenidae). Frontiers in zoology 2007, 4, 8.

(39) Rubinoff, D.; Cameron, S.; Will, K. A genomic perspective on the shortcomings of mitochondrial DNA for “barcoding” identification. Journal of heredity 2006, 97, 581– 594.

(40) Oord, A. v. d.; Li, Y.; Vinyals, O. Representation learning with contrastive predictive coding. arXiv preprint 1807.03748 2018,

(41) Hadsell, R.; Chopra, S.; LeCun, Y. Dimensionality reduction by learning an invariant mapping. 2006 IEEE computer society conference on computer vision and pattern recognition (CVPR’06). 2006; pp 1735–1742.

(42) Fabbri, A. R.; Kryściński, W.; McCann, B.; Xiong, C.; Socher, R.; Radev, D. Summeval: Re-evaluating summarization evaluation. Transactions of the Association for Computational Linguistics 2021, 9, 391–409.

(43) Robinson, J.; Sun, L.; Yu, K.; Batmanghelich, K.; Jegelka, S.; Sra, S. Can contrastive learning avoid shortcut solutions? Advances in neural information processing systems 2021, 34, 4974–4986.

(44) Paszke, A.; Gross, S.; Massa, F.; Lerer, A.; Bradbury, J.; Chanan, G.; Killeen, T.; Lin, Z.; Gimelshein, N.; Antiga, L.; others Pytorch: An imperative style, high-performance deep learning library. Advances in neural information processing systems 2019, 32.

(45) Portes, J.; Trott, A.; Havens, S.; King, D.; Venigalla, A.; Nadeem, M.; Sardana, N.; Khudia, D.; Frankle, J. MosaicBERT: A bidirectional encoder optimized for fast pretraining. Advances in Neural Information Processing Systems 2023, 36, 3106–3130.

(46) Dao, T. Flashattention-2: Faster attention with better parallelism and work partitioning. arXiv preprint 2307.08691 2023,

(47) Shazeer, N. Glu variants improve transformer. arXiv preprint 2002.05202 2020,

(48) Zhao, Q.; Zhang, C.; Zhang, W. dnaGrinder: a lightweight and high-capacity genomic foundation model. arXiv preprint 2409.15697 2024,

(49) Dao, T.; Fu, D.; Ermon, S.; Rudra, A.; Ré, C. Flashattention: Fast and memory-efficient exact attention with io-awareness. Advances in neural information processing systems 2022, 35, 16344–16359.

(50) Dauphin, Y. N.; Fan, A.; Auli, M.; Grangier, D. Language modeling with gated convolutional networks. International conference on machine learning. 2017; pp 933–941.

(51) Loshchilov, I.; Hutter, F. Decoupled weight decay regularization. arXiv preprint 1711.05101 2017,

(52) Akiba, T.; Sano, S.; Yanase, T.; Ohta, T.; Koyama, M. Optuna: A next-generation hyperparameter optimization framework. Proceedings of the 25th ACM SIGKDD international conference on knowledge discovery & data mining. 2019; pp 2623–2631.

(53) Dalla-Torre, H.; Gonzalez, L.; Mendoza-Revilla, J.; Lopez Carranza, N.; Grzywaczewski, A. H.; Oteri, F.; Dallago, C.; Trop, E.; de Almeida, B. P.; Sirelkhatim, H.; others Nucleotide Transformer: building and evaluating robust foundation models for human genomics. Nature Methods 2025, 22, 287–297.

(54) Nguyen, E.; Poli, M.; Faizi, M.; Thomas, A.; Wornow, M.; Birch-Sykes, C.; Massaroli, S.; Patel, A.; Rabideau, C.; Bengio, Y.; others Hyenadna: Long-range genomic sequence modeling at single nucleotide resolution. Advances in neural information processing systems 2023, 36, 43177–43201.

(55) McInnes, L.; Healy, J.; Melville, J. Umap: Uniform manifold approximation and projection for dimension reduction. arXiv preprint 1802.03426 2018,

(56) Everitt, B. S.; Landau, S.; Leese, M.; Stahl, D.; others Hierarchical clustering. Cluster analysis 2011, 5, 71–110.

(57) Cover, T.; Hart, P. Nearest neighbor pattern classification. IEEE transactions on information theory 1967, 13, 21–27.

(58) Karollus, A.; Hingerl, J.; Gankin, D.; Grosshauser, M.; Klemon, K.; Gagneur, J. Species-aware DNA language models capture regulatory elements and their evolution. Genome biology 2024, 25, 83.

(59) Wang, T.; Isola, P. Understanding contrastive representation learning through alignment and uniformity on the hypersphere. International conference on machine learning. 2020; pp 9929–9939.

(60) Peterson, J.; Garges, S.; Giovanni, M.; McInnes, P.; Wang, L.; Schloss, J. A.; Bonazzi, V.; McEwen, J. E.; Wetterstrand, K. A.; Deal, C.; others The NIH human microbiome project. Genome research 2009, 19, 2317–2323.

(61) Yang, H.; Cole, J.; Li, K. Automating Large-scale In-silico Benchmarking for Genomic Foundation Models. arXiv preprint 2410.01784

(62) Lee, N. K.; Tang, Z.; Toneyan, S.; Koo, P. K. EvoAug: improving generalization and interpretability of genomic deep neural networks with evolution-inspired data augmentations. Genome Biology 2023, 24, 105.

(63) Zhang, Y.; Zhu, R.; Zhang, S.; Zhou, X.; Chen, S.; Chen, X. Feature Augmentation for Self-supervised Contrastive Learning: A Closer Look. 2024 International Joint Conference on Neural Networks (IJCNN). 2024; pp 1–8.

(64) Cao, B.; Li, X.; Wang, B.; He, T.; Zheng, Y.; Zhang, X.; Zhang, Q. Achieving handle-level random access in an encrypted DNA archival storage system via frequency dictionary mapping coding. Patterns 2025,

(65) Cao, B.; Zhao, Y.; Lei, X.; Shao, Q.; Wang, K.; Wang, B.; Zhou, S.; Pan, Z. DBSP: An End-to-end Pipeline for DNA Storage Data Reconstruction from DNA Sequencing. IEEE Transactions on Molecular, Biological, and Multi-Scale Communications 2025, 1, 1.

(66) Rasool, A.; Hong, J.; Hong, Z.; Li, Y.; Zou, C.; Chen, H.; Qu, Q.; Wang, Y.; Jiang, Q.; Huang, X.; others An Effective DNA-Based File Storage System for Practical Archiving and Retrieval of Medical MRI Data. Small Methods 2024, 8, 2301585.

(67) Rasool, A.; Jiang, Q.; Wang, Y.; Huang, X.; Qu, Q.; Dai, J. Evolutionary approach to construct robust codes for DNA-based data storage. Frontiers in Genetics 2023, 14, 1158337.

(68) Rasool, A. RFS-codec: A Novel Encoding Approach to Store Image Data in DNA. Journal of Artificial Intelligence in Bioinformatics 2025, 1, 41–50.

